# Disruption of RBM20 causes atrial electrophysiological disturbances

**DOI:** 10.64898/2026.03.14.711772

**Authors:** Leo Weirauch, Felix Wiedmann, Laura Schraft, Maarten M.G. van den Hoogenhof, Merten Prüser, Manuel Kraft, Yinuo Wang, Amelie Paasche, Gergana Dobreva, Lars Steinmetz, Constanze Schmidt

**Affiliations:** Department of Cardiology, University Hospital Heidelberg, Heidelberg, Germany; DZHK (German Center for Cardiovascular Research), partner site Heidelberg /Mannheim, University of Heidelberg, Heidelberg, Germany; Department of Cardiology and Pneumology, University Medical Center Goettingen, Goettingen, Germany; DZHK (German Center for Cardiovascular Research), partner site Lower Saxony, University of Goettingen, Goettingen, Germany; European Molecular Biology Laboratory (EMBL), Genome Biology Unit, Heidelberg, Germany; Institute of Experimental Cardiology, University Hospital Heidelberg, Heidelberg, Germany; Department of Cardiovascular Genomics and Epigenomics, European Center for Angioscience (ECAS), Medical Faculty Mannheim, Heidelberg University, Mannheim, Germany; Department of Genetics, Stanford University School of Medicine, Stanford, CA, USA

**Keywords:** Atrial cardiomyopathy, Atrial fibrillation, Dilated cardiomyopathy, Electrophysiological remodeling, *RBM20* mutation, SGLT inhibitors

## Abstract

**Background:** Dilated cardiomyopathy (DCM) is a leading cause of heart failure, with 30–50 % of cases attributed to familial inheritance. Mutations in RNA-binding motif protein 20 (RBM20) account for 3–5 % of cases and are associated with severe DCM and ventricular arrhythmias. However, the role of *RBM20* mutations in atrial cardiomyopathy (AtCM) and atrial fibrillation (AF) remains underexplored. This study investigates the effects of the *RBM20*-R636Q mutation on atrial electrophysiology and evaluates sodium-glucose co-transporter (SGLT) inhibitors as potential therapeutics.

**Results:** *Rbm20*-R636Q mice exhibited atrial remodeling, including hypertrophy, left atrial enlargement, and shortened action potential duration at 90% repolarization (APD_90_). Compared with *RBM20*-knockout and laminopathy models, *RBM20*-R636Q mice showed distinct reductions in *I*_to_ / *I*_Kur_ without changes in *I*_K,sus_ or *I*_K,tail_ currents, alongside TASK-1 potassium current upregulation and alterations of *I*_CaL_. SGLT inhibitors (sotagliflozin, empagliflozin, dapagliflozin) reduced AP inducibility and partially restored APD_90_, with effects comparable to lidocaine, suggesting a role in modulating peak sodium currents.

**Conclusions:** *RBM20* mutations contribute to atrial remodeling, promoting AtCM and AF. SGLT inhibitors demonstrate therapeutic potential by modulating atrial electrophysiology and reducing arrhythmogenesis, offering a promising strategy for managing RBM20-related cardiac disorders.

**Graphical Abstract:** 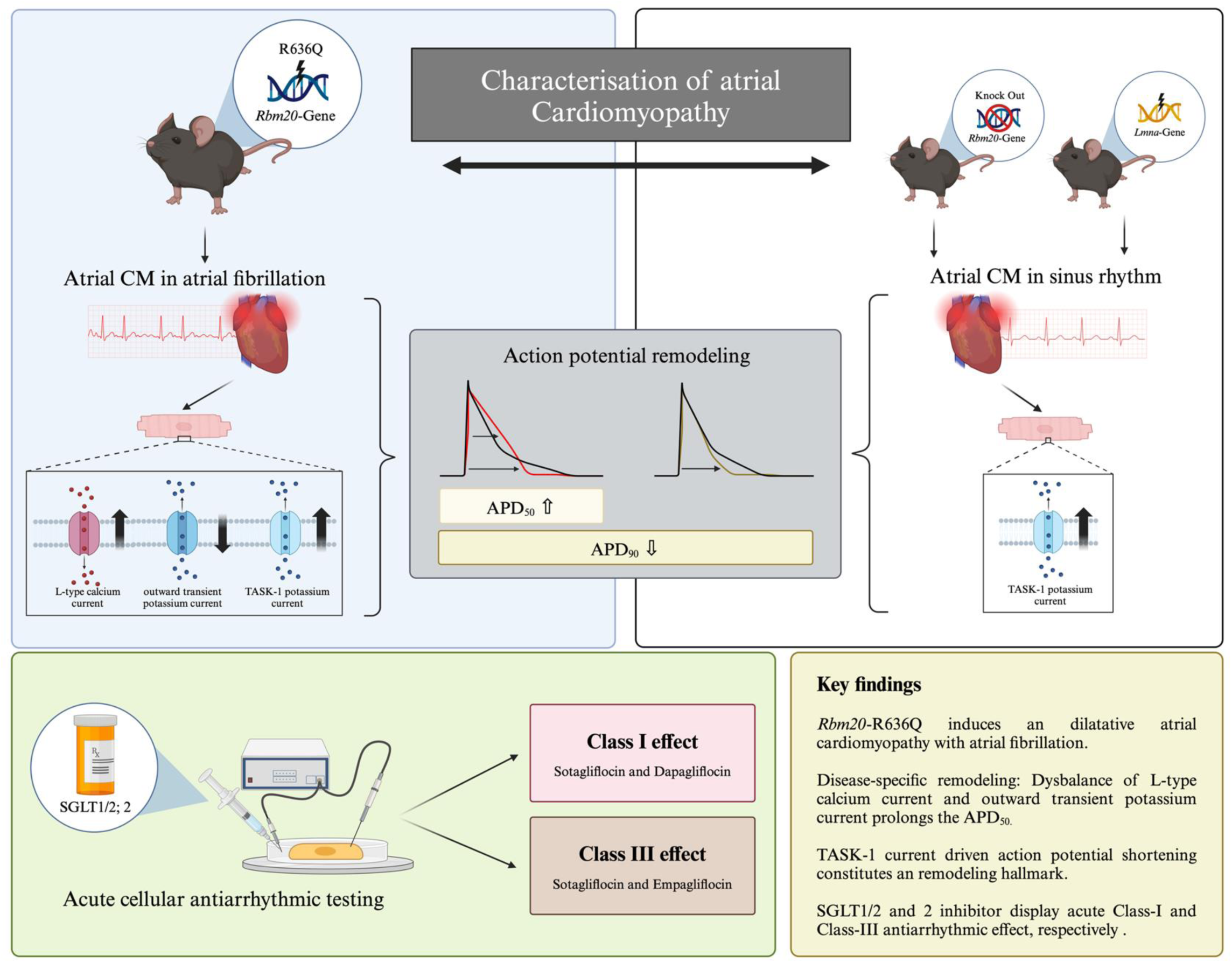

## Introduction

Splice-regulatory networks play a pivotal role in generating transcriptomic diversity, contributing to the functional specialization of different cell types across organ systems^1^. Among these regulators, RNA-binding motif protein 20 (RBM20) is a crucial orchestrator of cardiac isoform expression, influencing the splicing of over 30 essential targets involved in sarcomere structure, ion handling, and calcium signaling^2–4^. Mutations in *RBM20* are associated with a clinically aggressive form of dilated cardiomyopathy (DCM), characterized by progressive left ventricular dysfunction, heart failure (HF), and a high incidence of life-threatening arrhythmias^3,5^. There is growing evidence to suggest that arrhythmias in patients with *RBM20* mutations may not solely be a consequence of structural heart disease but could also arise from specific electrical dysfunctions^4,6,7^. For instance, the RBM20’s regulatory impact on critical ion-handling proteins, such as ryanodine receptor 2 and Calcium/calmodulin-dependent protein kinase type II delta (CaMKIIδ), plays a fundamental role in cardiac electrophysiology, particularly with regard to arrhythmogenesis^5,8^.

The rising global prevalence of atrial fibrillation (AF) has sparked renewed need in understanding the molecular background of atrial cardiomyopathy (AtCM), underlying atrial arrhythmogenesis^9,10^. A recently published clinical consensus statement on AtCM has suggested that mutations in *RBM20* and *TTN* (titin), a known splicing target of RBM20, could contribute to atrial remodeling and arrhythmogenesis, potentially independent of left ventricular dysfunction, underscoring the need to investigate the role of RBM20 in AtCM and AF more deeply^10^. Notably, genome-wide association studies (GWAS) have linked single nucleotide polymorphisms (SNP) in both, the genomic region encoding for RBM20 and its splicing target titin to early-onset AF^11,12^. This association has been further recognized by the 2023 ACC/AHA/ACCP/HRS Guideline for the Diagnosis and Management of AF, which now include a class IIb recommendation for genetic testing in AF patients aged 45 years or younger, with both *TTN* and *RBM20* among the key genes potentially implicated^13,14^.

Although the role of *RBM20* mutations in ventricular dysfunction and arrhythmias is relatively well-established, much remains unknown about the specific molecular mechanisms linking these splicing defects to atrial arrhythmias. In *Rbm20*-knockout (KO) mouse models, altered calcium handling has been identified as a critical factor contributing to increased susceptibility to arrhythmias^8,15^. However, these findings have primarily focused on ventricular cardiomyopathy, leaving the contribution of atrial remodeling and its link to AF underexplored. Recent studies on Rbm20 dysfunction have demonstrated that mice and rats exhibit severe atrial remodeling with frequent development of AF, even in the absence of significant left ventricular dysfunction^16,17^. Further, the results of different clinical studies indicate that *RBM20* mutations affecting the RS domain, which is a critical region for splicing activity, are associated with a markedly elevated prevalence of AF^16,18^. These variants were shown to mislocalize to the cytoplasm, leading to formation of detrimental ribonucleoprotein granules, which has been shown to worsen the left ventricular function in animals ^17,19^. Again, these mutations seem to predispose patients to atrial arrhythmias without exacerbating left ventricular function to the same extent as other *RBM20* variants, underscoring the importance of cardiac electrophysiology in these specific mutations^16^.

Mechanism-based therapeutic interventions for *RBM20*-associated DCM remain limited, despite early translational studies demonstrating promising outcomes with CRISPR-based gene therapy approaches^20,21^. Sodium-glucose cotransporter inhibitors (SGLTi), initially developed as oral antidiabetic agents, have recently shown efficacy in reducing adverse events in HF patients with both reduced and preserved left ventricular function, independent of glycemic control^22,23^. Moreover, clinical studies have documented a decreased incidence of AF following SGLTi treatment^24–26^, with preliminary molecular investigations attributing this effect to direct effects on the cardiac actions potential (AP)^27^.

Given the rising recognition of *RBM20’s* role in AtCM, this study aims to characterize the cellular electrophysiological remodeling patterns associated with *RBM20* mutations on a transcriptional and cellular electrophysiological level, revealing key ion channel alterations contributing to atrial arrhythmogenesis. Additionally, SGLT inhibitors were evaluated as potential antiarrhythmic therapies, demonstrating promising effects on action potential morphology in *RBM20*-mutant cardiomyocytes (see Graphical Abstract).

## Methods

A detailed description of all experimental procedures is provided in the Supplementary Material.

### Animal ethics and mouse models

All animal procedures were approved by the local Animal Welfare Committee (Regierungspräsidium Karlsruhe; G233/17, G225/20, G-194/18) and conducted in accordance with German and EU animal protection laws, NIH guidelines (NIH Pub. No. 86-23), and EMBL institutional policies. *Rbm20-R636Q* knock-in mice were generated by zygotic microinjection of Cas9 and crRNA:tracrRNA complexes targeting *Rbm20*, using a single-stranded DNA donor template^20^. *Rbm20* knockout (KO) mice were initially generated at the Academic Medical Center, Amsterdam, using a conditional allele and Cre-mediated recombination, as previously described^8,28^. The *Lmna* tm1.1Yxz/J line, was obtained from Jackson Laboratory and maintained on a C57BL/6J background as reported^29,30^

### Animal handling and phenotyping

Mice were housed under standard conditions with ad libitum access to food and water. Experiments were performed on male and female mice aged 15–17 weeks unless stated otherwise. ECG recordings were performed in conscious mice using the ecgTUNNEL system (emka Technologies), allowing non-invasive, longitudinal 6-lead monitoring. Data were acquired using ecgAUTO software and corrected for heart rate using Fridericia’s formula^31^.

### Cardiomyocyte isolation and electrophysiology

Atrial cardiomyocytes were enzymatically isolated post-euthanasia using collagenase and protease digestion. Cells were maintained in tailored cardioplegic and storage solutions before electrophysiological analysis. Whole-cell patch-clamp recordings were conducted using borosilicate pipettes and an Axopatch 200B amplifier (Molecular Devices). Action potentials (APs) were recorded under current clamp, while L-type Ca²⁺, Na⁺, and K⁺ currents were obtained via voltage-clamp protocols using established solutions and stimulation paradigms, as described previously^32–34^.

### Transcriptomics

RNA from atrial tissue was extracted using TRIzol and subjected to poly(A)-enriched RNA-seq (Illumina NextSeq 2000). Reads were aligned to GRCm39 using STAR^35^, quantified with featureCounts^36^, and analyzed for differential expression with DESeq2^37^. Genes with adjusted *p* < 0.01 and |log₂ fold change| > 0.5 were considered differentially expressed.

## Results

Clinical studies have linked mutations in the RSRSP domain of RBM20, a critical region nuclear localization of the protein, as being associated with a high prevalence of AF^17^. The *Rbm20*-R636Q mutation in mice, which is located within this RSRSP stretch, serves as an ortholog to the *RBM20-*R634Q mutation in humans, previously found in DCM patients. While previous investigations have comprehensively characterized the *Rbm20*-R636Q mice in terms of the molecular splicing defect, cytoplasmic mislocalization, ventricular DCM phenotype, and overall survival, we aimed to specifically explore the atrial phenotype of these mice^3,20^. The R636Q mutation was selected due to its localization within the critical RBM20 RSRSP domain, which is associated with significant arrhythmogenic potential, thus allowing exploration of atrial electrophysiological remodeling independent of the previously established ventricular phenotype.

### Rbm20-R636Q knock-in mice exhibit an AtCM phenotype with spontaneous episodes of AF

To confirm the cardiomyopathy phenotype at the atrial level, echocardiography was conducted, and the LA area was quantified. In homozygous *Rbm20*-R636Q knock-in mice, a nearly two-fold increase in LA area was observed compared to wild-type (wt) controls (p = 0.0012; n = 10), measured in the parasternal long-axis view (Fig. 1a). At the cellular level, this enlargement was accompanied by significant hypertrophy of atrial cardiomyocytes. When membrane capacitance was measured as a surrogate for cell size in whole-cell patch-clamp recordings, a 29.2 ± 9.5 % increase in cell size was found in *Rbm20*-R636Q mice compared to wild-type (p = 0.036; n/N = 21/10; Fig. 1b).

**Figure 1:**
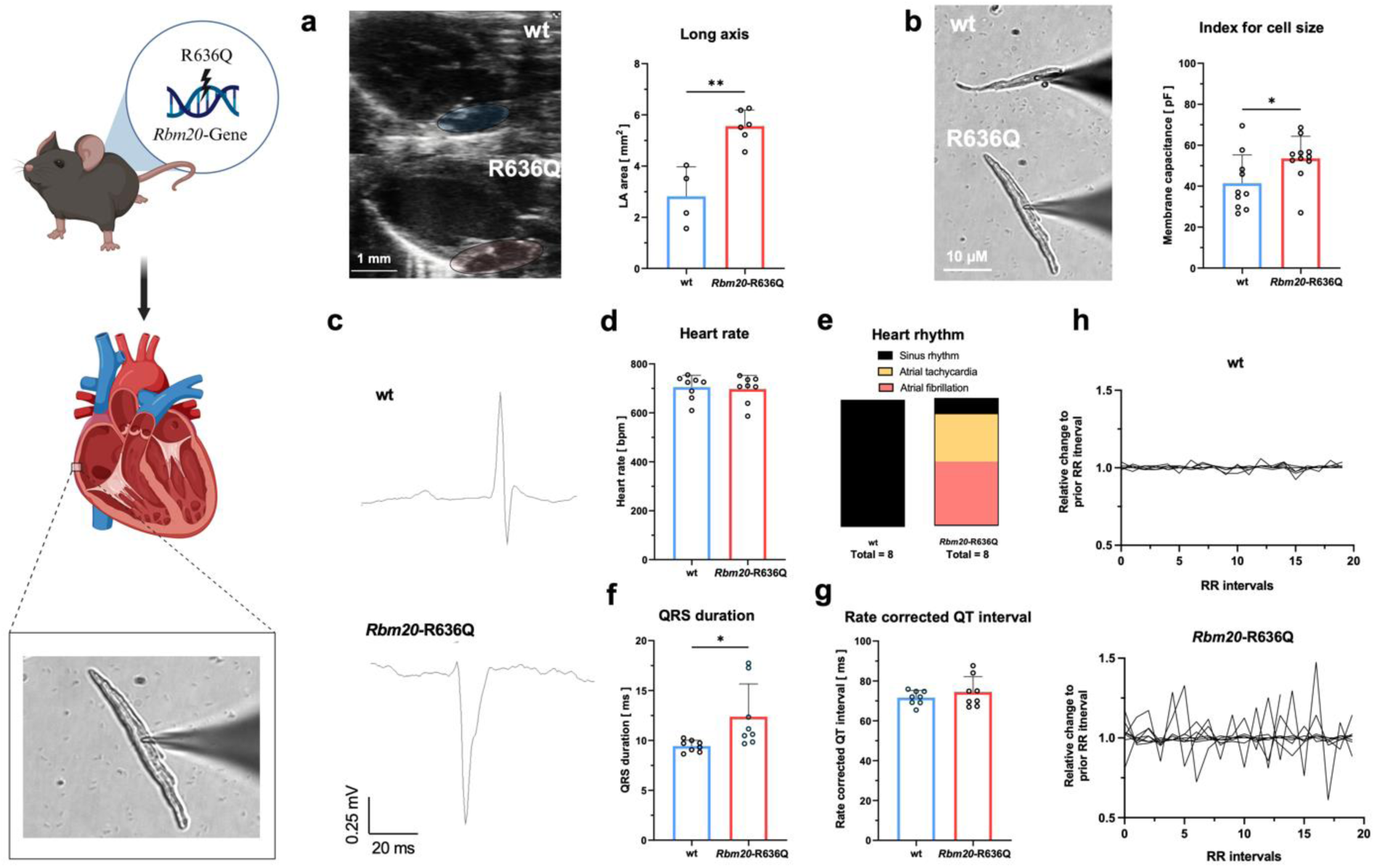
Homozygous Rbm20-R636Q knock-in mice exhibit structural and electrophysiological features of atrial cardiomyopathy. *Left:* Schematic illustration of the experimental protocol from organ harvest until cardiomyocyte isolation **a** Representative echocardiographic images obtained from the parasternal long-axis view in wild-type (wt, n = 4) and homozygous *Rbm20*-R636Q knock-in mice (n = 6), with corresponding quantification of the left atrial (LA) area (scalebar: 1 mm, as denoted). **b** Representative images of atrial cardiomyocytes, isolated from wt and *Rbm20*-R636Q mice attached to patch-clamp pipettes are presented on the left-hand side (scalebar: 10 µm). *Right:* Membrane capacitances, which serve as an index of cell size, are compared between atrial cardiomyocytes isolated from wt (n = 10 cells of N = 5 individuals) and *Rbm20*-R636Q (n/N = 11/5) mice. **c** Representative surface ECG traces, recorded from wt or *Rbm20*-R636Q mice. **d** Mean heart rate (averaged over 1–2 min), **e** underlying rhythm, **f** QRS duration and **g** rate corrected QT interval (using the Fridericia formula) of wt or *Rbm20*-R636Q mice (n = 8 each). **h** Relative variation in 20 representative R-R intervals, with each line representing one individual animal pointing towards an increased incidence of atrial fibrillation in *Rbm20*-R636Q mice. All data are presented as mean ± SEM, scalebars are provided as insets. p values were calculated using two-tailed unpaired Student’s t-tests (*, p < 0.05; **, p < 0.01).

To further characterize the atrial electrophysiological phenotype, non-invasive ECG measurements were performed using the emka ecgTUNNEL system. Representative surface ECGs are shown in Fig. 1c. Both groups showed similar heart rates (p = 0.77; n = 16; Fig. 1d); however, three out of eight *Rbm20*-R636Q mice exhibited episodes of atrial tachycardia, as characterized by a lack of P waves in all surface ECG leads and an increased atrial frequency. Four out of eight mice also showed an irregular ventricular excitation which then was defined as AF as underlying heart rhythm. (Fig. 1c, e; Suppl. Fig. S2).

Moreover, the QRS duration, as an indicator of ventricular cardiomyopathy, was significantly prolonged by 23.81 ± 9.5 % in *Rbm20*-R636Q mice (p = 0.025; n=16; Fig. 1f). No significant differences were detected with respect to the heart rate-corrected QT interval (p = 0.37; n=16; Fig. 1g). Finally, the relative variation in 20 representative R-R intervals was higher in the *Rbm20*-R636Q group, again reflecting the presence of AF (Fig. 1h). Interestingly after a subdivision according to gender, the significant increase of QRS duration was present in male *Rbm20*-R636Q mice only (p = 0.0372; n =3-4; Suppl. Fig. S3). As previously described, the echocardiographically measured left ventricular ejection fraction was moderately to severely reduced in *Rbm20*-*R636Q* mice (Suppl. Fig. S8). This was accompanied by a significant increase in left ventricular end-diastolic and end-systolic volumes, as well as a significant decrease in left ventricular longitudinal strain (Suppl. Fig. S8). Furthermore, the echocardiographically quantified left atrial ejection fraction (LAEF) and the left atrial reservoir strain showed significant reduction in Rbm20-R636Q mice, compared to WT controls (Suppl. Fig. S9). Interestingly, reduction of LAEF was significantly more pronounced, compared to reduction in LVEF (Suppl. Fig. S9).

### The Rbm20-R636Q mutation causes alterations in cellular electrophysiology of isolated atrial cardiomyocytes

Given the established link between mutations in the *RBM20* gene and cardiac electrical disturbances, the electrophysiological properties of isolated atrial cardiomyocytes were assessed using patch-clamp measurements. APs recorded from *Rbm20*-R636Q mutant atrial cardiomyocyte (CMs) displayed notable morphological changes compared to wt-cells (Fig. 2a). To define those alterations the AP duration is quantified at the timepoints of 50 % and 90 % repolarization as indicated in Fig. 2a, respectively. Specifically, the AP duration at 50 % repolarization (APD_50_) was significantly prolonged by 40.0 ± 15.5 % (p = 0.021; n/N = 29/10), whereas the AP duration at 90 % repolarization (APD_90_) exhibited a 29.7 ± 7.6 % shortening (p = 0.0011; n = 29/10; Fig. 2a). Despite these changes, the maximum upstroke velocity and AP amplitude remained unaltered, resulting in a triangular AP morphology, a characteristic feature of functional atrial electrical remodeling in AtCM. Importantly, these alterations were found to be present in left and right atrial CMs and in both male and female mice (Suppl. Fig. S4–5), except the observation of QRS prolongation which was only present in male mice. Further, no significant differences were observed when comparing young mice as the age of 16 ± 1 weeks (Fig. 2) with older ones (30± 1 weeks; Suppl. Fig. S6).

**Figure 2:**
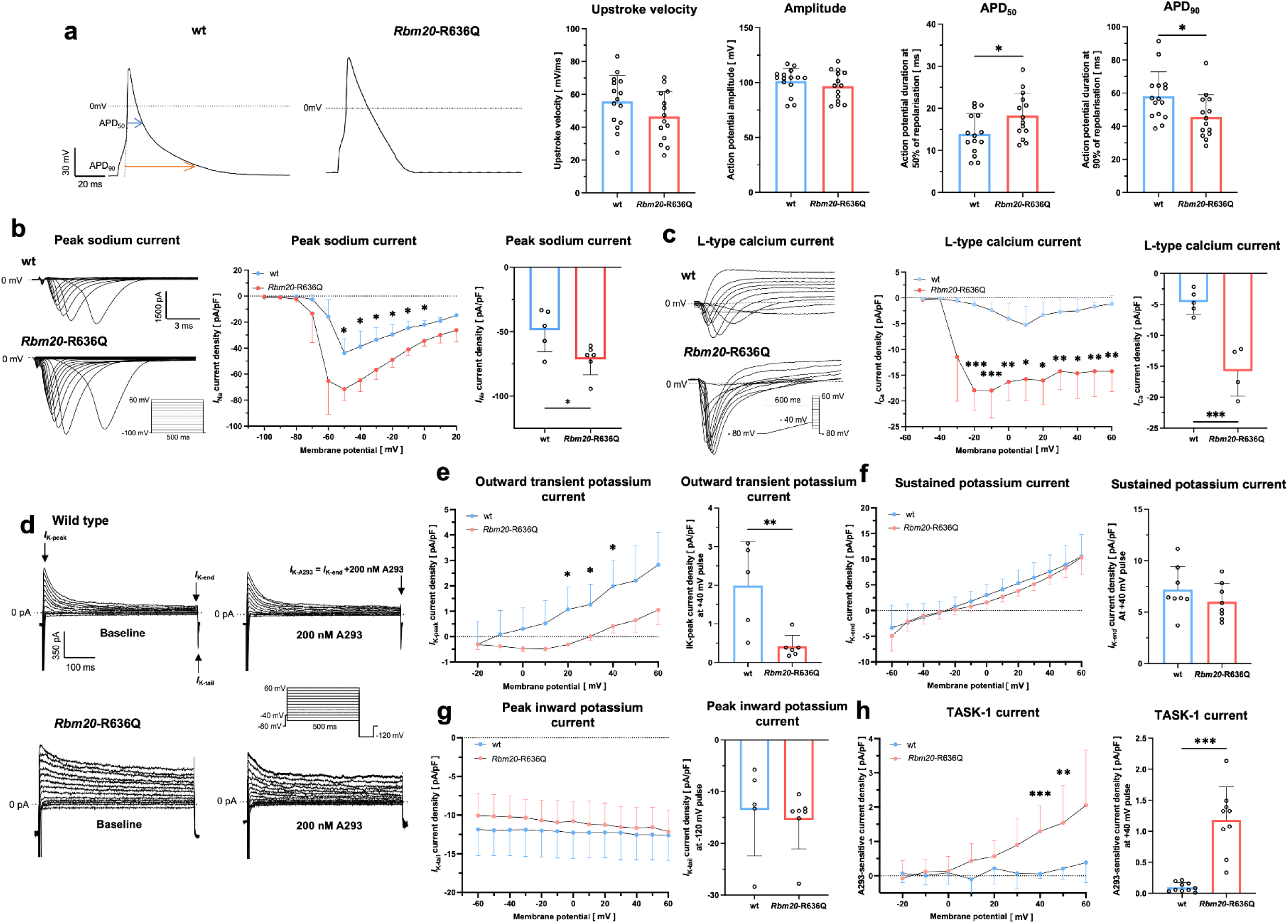
Cellular electrical remodeling in atrial cardiomyocytes, isolated from Rbm20-R636Q knock-in mice. **a** *Left:* Representative action potentials (APs), recorded from single atrial cardiomyocytes (CM), isolated from wild-type (wt) and homozygous *Rbm*20-R636Q knock-in mice using the patch-clamp technique in current clamp configuration. *Right*: Comparison of maximum upstroke velocity, AP amplitude, and AP duration at 50% or 90% of repolarization among wt (n = 15 cells from N = 6 animals) and *Rbm*20-R636Q (n/N = 14/5) mice. **b** *Left*: Representative peak sodium currents, recorded from wt and *Rbm20*-R636Q CMs using patch-clamp measurements in the voltage clamp configuration by application of the pulse protocol depicted. *Center*: comparison of sodium current-voltage (I-V) relationships from wt (n/N = 5/3) and *Rbm20*-R636Q (n/N = 6/3) CMs. *Right*: peak sodium current densities quantified at −50 mV. **c** *Left*: Representative L-type calcium currents, recorded from wt and *Rbm20*-R636Q CMs using patch-clamp measurements in the voltage clamp configuration by application of the pulse protocol depicted. *Center*: comparison of L-type calcium current-voltage (I-V) relationships from wt (n/N = 5/4) and *Rbm20*-R636Q (n/N = 4/4) CMs. *Right*: L-type calcium densities quantified at −10 mV. **d** Representative potassium currents recorded from wt and *Rbm20*-R636Q CMs using patch-clamp measurements in the voltage clamp configuration before and after application of the TASK-1 inhibitor A293 (200 nM), by application of the pulse protocol depicted. **e** Comparison of *I*_K,Peak_ (equivalent to transient outward potassium currents *I*_to_ and the fast-activating slow-inactivating K^+^ current *I*_K,slow_) current-voltage (I-V) relationships and mean current densities quantified at +40 mV from wt (n/N = 5/3) and *Rbm20*-R636Q (n/N = 6/3) CMs. **f** Comparison of current-voltage (I-V) relationships from the sustained potassium current component *I*_K,sus_ and mean current densities quantified at +40 mV from wt (n/N = 8/3) and *Rbm20*-R636Q (n/N = 8/3) CMs. **g** Voltage-dependence of *I*_K,Tail_ (equivalent to peak inward potassium current) and mean current densities, quantified at −120 mV from wt (n/N = 5/3) and *Rbm20*-R636Q (n/N = 7/3) CMs. **h** Comparison of TASK-1 current-voltage (I-V) relationships (*I*_TASK-1_), difference to baseline quantified after the application (10 min.) of the high-affinity TASK-1 inhibitor A293 (200 nM). Mean current densities are quantified at +40 mV from wt (n/N = 10/4) and *Rbm20*-R636Q (n/N = 9/4) CMs. All data are presented as mean ± SEM. Dashed lines represent zero current /potential levels. Scalebars and pulse protocols are provided as insets, p values were determined using two-tailed unpaired Student’s t-tests (*, p < 0.05; **, p < 0.01; ***,p < 0.001; ***,p < 0.001).

To gain further insight into the underlying molecular mechanisms of these AP changes, key currents that form the cardiac APs were compared in isolated atrial CMs of both experimental groups. In *Rbm20*-R636Q CMs, peak sodium currents responsible for the AP upstroke were significantly increased by 47.9 ± 16.3 % (p = 0.01; n/N = 11/6; Fig. 2b).

Next, the activity of L-type calcium channels, which play a key role in the plateau phase of the cardiac AP, was compared among atrial CM, isolated from mutant and wt mice. Here, a 3-fold increase in L-type calcium current density (p = 0.023; n/N = 9/6; quantified at +10 mV) was found in *Rbm20*-R636Q mice, indicating enhanced Ca^2+^ influx during the plateau and early repolarization phase (Fig. 2c).

Lastly, potassium (K^+^) currents, critical for AP repolarization, were examined. The outward transient K^+^ current (*I*_K,peak_), which comprises the transient outward (*I*_to_) as well as the ultra-rapid delayed rectifier current components (*I*_Kur_) was significantly reduced by 79.8 % in *Rbm20*-R636Q CMs (p = 0.0092; n = 11–16/6; Fig. 2d–e). With respect to the peak inward K^+^ current (*I*_K,tail_) and the sustained K^+^ current (*I*_K,sus_), however, no alternations were observed (Fig. 2d, f, g). Moreover, we assessed the atrial TASK-1 potassium current, a key contributor to atrial electrical remodeling^33,34,38^. Selective inhibition of TASK-1 with the specific inhibitor A293 (200 nM) revealed a 12-fold increase in TASK-1 current density in *Rbm20*-R636Q CMs compared to wt (p < 0.0001; n = 19/8; Fig. 2h). This increase likely contributes to the shortening of APD_90_ and the elevated susceptibility to atrial arrhythmias observed in these animals. All together, these data link the AP abnormalities to the respective ion currents underlying them.

### Electrophysiological changes in a murine Rbm20-KO model

To further investigate the mechanisms underlying the AtCM phenotype in *Rbm20*-R636Q mice, comparisons were made with a murine *Rbm20*-KO model. Using the emka ECG-Tunnel system, it was observed that the mean heart rate in *Rbm20*-KO mice was significantly lower compared with wt (19.3 ± 7.2 %; p = 0.016; n = 8; Fig. 3a). While the P-wave duration remained unchanged, the P-wave amplitude was significantly increased by 16.8 ± 5.5 % in *Rbm20*-KO mice compared to wild-type (p = 0.019; n = 9; Fig. 3a). No significant differences were detected in the PR interval, QRS duration, or rate-corrected QT interval (Fig. 3a).

**Figure 3:**
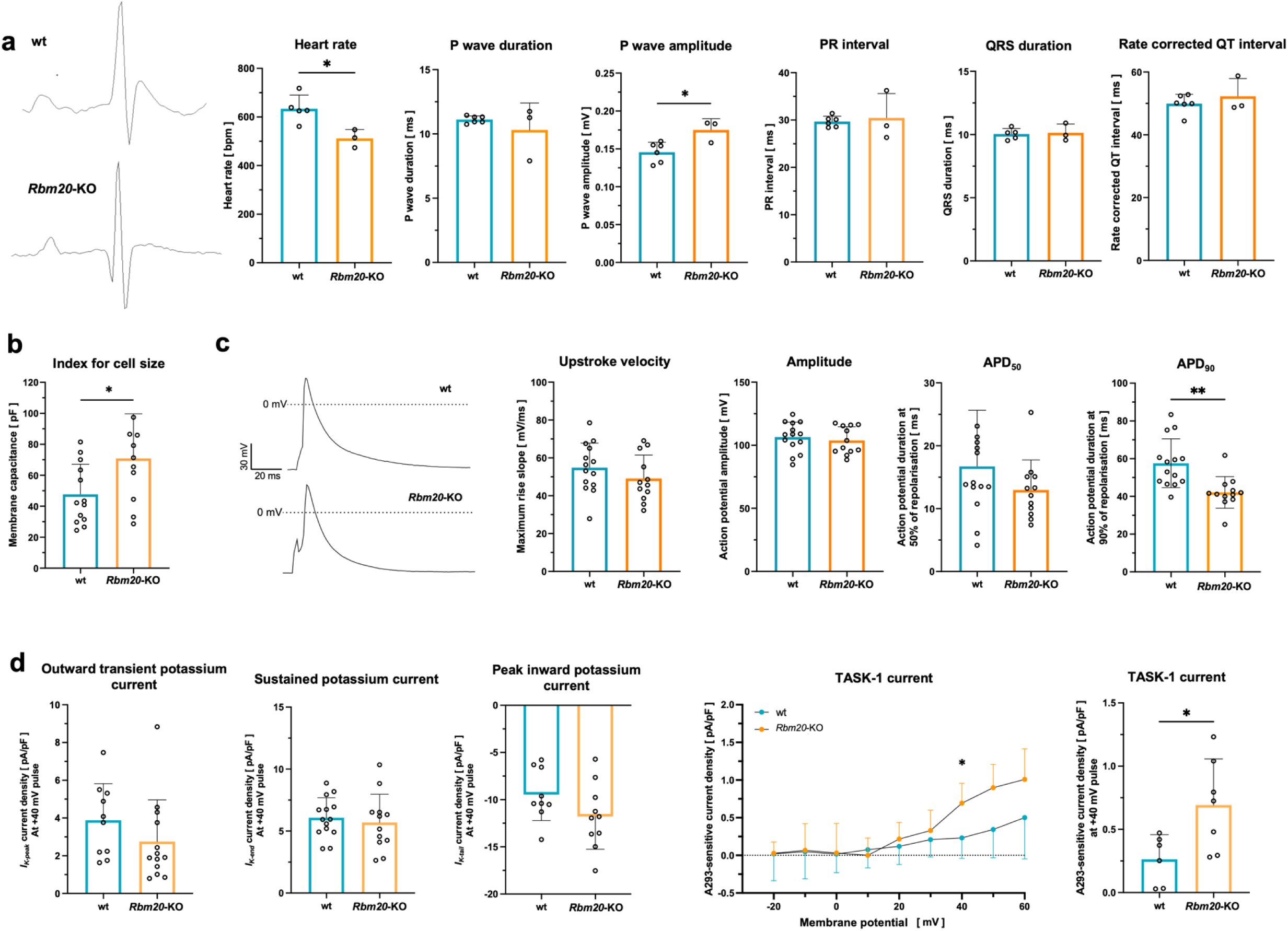
Electrophysiological characterization of Rbm20-knockout mice. **a** Representative surface ECG traces, recorded from wild-type (wt) and *Rbm20*-knockout (KO) mice. Comparison of mean heart rate (averaged over 1–2 min), P-wave duration, P-wave amplitude, PR-interval, QRS-duration and rate corrected QT interval (using the Fridericia formula) of wt (n = 5–6) or *Rbm20*-KO mice (n = 3). **b** Membrane capacitances, which serve as an index of cell size, are compared between atrial cardiomyocytes isolated from wt (n = 13 cells of N = 4 individuals) and *Rbm20*-KO (n/N = 10/5) mice. **c** Representative action potentials (APs), recorded from single atrial cardiomyocytes (CM) of both animal groups using the patch-clamp technique in current clamp configuration. Comparison of maximum upstroke velocity, AP amplitude, and AP duration at 50% or 90% of repolarization among wt (n/N = 13–14/4 animals) and *Rbm*20-KO (n/N = 12/4) mice. **d** Comparison of mean current densities of *I*_K,Peak_ (equivalent to ultra rapid and transient outward potassium currents *I*_Kur_ and *I*_to_), quantified at +40 mV, mean current densities of the sustained potassium current component *I*_K,sus_ quantified at +40 mV, *I*_K,Tail_ (equivalent to peak inward potassium current) and TASK-1 current-voltage (I-V) relationships, measured by application of the high-affinity TASK-1 inhibitor A293 (200 nM). Mean current *I*_TASK-1_ densities are quantified at +40 mV. Mean current values± SEM, derived from wt (n/N = 6–14/4) and *Rbm20*-KO (n/N = 7–15/4) CMs are displayed for each potassium current component. Data are presented as mean ± SEM. Dashed lines represent zero potential levels. Scalebars are provided as insets, p values were determined using two-tailed unpaired Student’s t-tests (*, p < 0.05; **, p < 0.01).

Atrial CMs hypertrophy was also observed in *Rbm20*-KO mice, with membrane capacitance (used as a surrogate for cell size) significantly increased by 48.7 ± 20.8% compared to wt (p = 0.029; n/N = 23/9; Fig. 3b). Current-clamp recordings of APs from isolated atrial CMs revealed no significant differences in maximum upstroke velocity, AP amplitude and APD_50_. However, APD_90_ was significantly shortened by 36.6 ± 10.3 % (p = 0.0016; n/N = 25-26/8;) (Fig. 3c). Overall, the *Rbm20*-KO mice exhibited a shortened and triangularized AP, though the exact pattern differed slightly from the *Rbm20*-R636Q mice.

Regarding the underlying ionic currents, no significant changes were observed in *I*_K,Peak_, *I*_K,sus_, or *I*_K,tail_ currents. However, a 2.6-fold upregulation of the TASK-1 current was again detected (p = 0.026; n/N = 13-29/8; Fig. 3d).

### Characterization of AtCM in a DCM disease model of Lmna^+/-^ laminopathy mice

To investigate whether the observed AtCM in *Rbm20*-R636Q mice arises as a secondary consequence of systolic HF within the context of DCM or is directly linked to atrial-specific effects of the *Rbm20*-R636Q mutation, a comparative analysis was conducted using an alternative murine DCM model. This *Lmna*^+/-^ mouse model, which induces a laminopathy-related DCM through haploinsufficiency of the *Lmna* gene is also characterized by severe left-ventricular dysfunction. While previous studies have thoroughly characterized the ventricular DCM phenotype and underlying molecular pathology in *Lmna*^+/-^ mice, the focus of this study was to assess the atrial cardiomyopathy phenotype in depth.

Compared to wt controls, *Lmna*^+/-^ mice exhibited comparable mean heart rates and P-wave durations measured at awake animals using the emka ECG-Tunnel system (Fig. 4a). However, a distinct atrial low-voltage pattern was observed in the laminopathy mice, with P-wave amplitudes reduced by 29.5 ± 10.8 % as compared to wt controls (p = 0.025; n = 12; Fig. 4a). Despite this evidence of atrial electrical remodeling, no episodes of spontaneous AF were detected in this model. Furthermore, no significant differences were noted with respect to the PR interval, QRS duration, or rate-corrected QT interval (n = 12; Fig. 4a).

**Figure 4:**
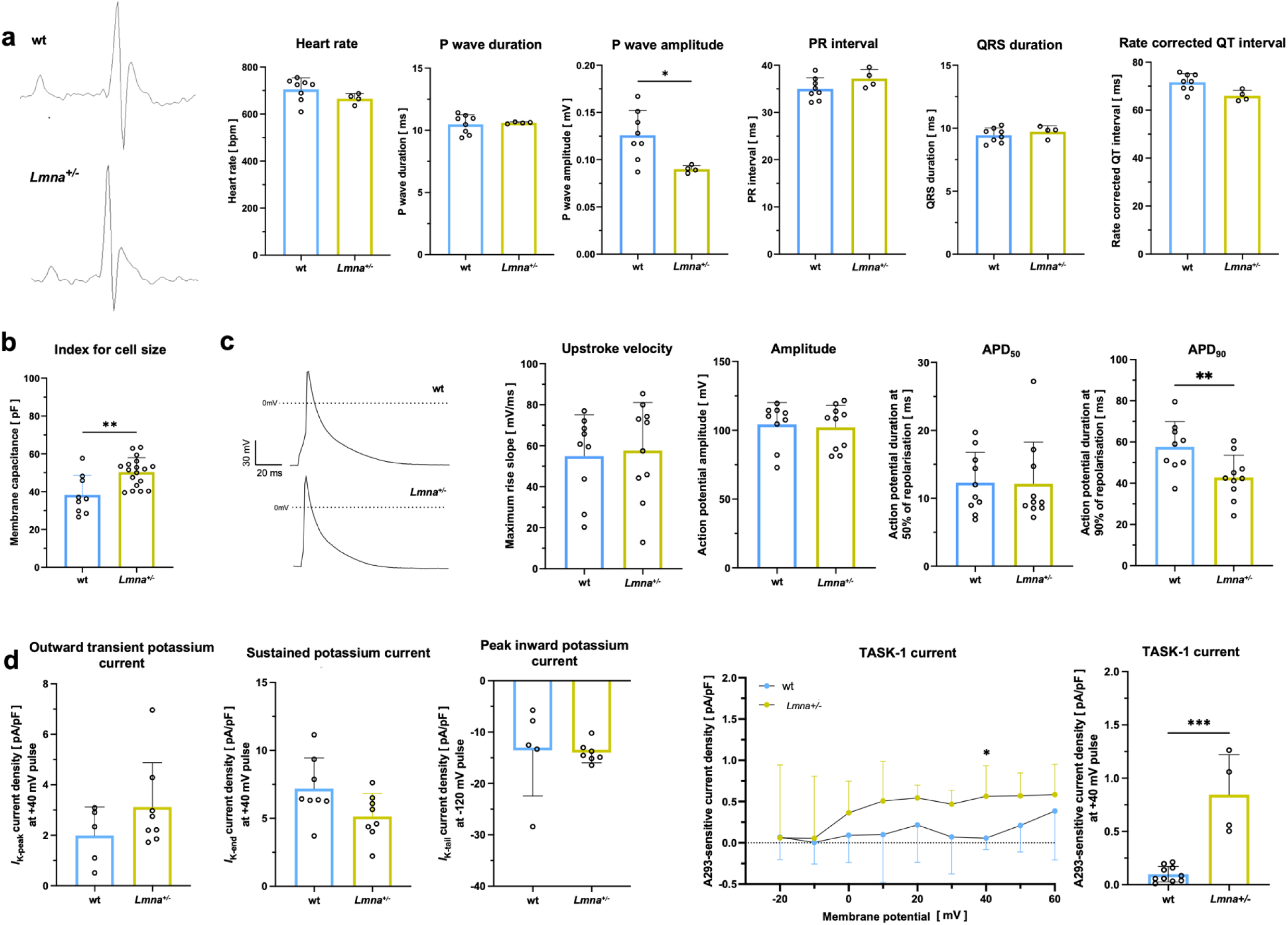
Electrophysiological characterization of Lmna^+/-^ mice. **a** Representative surface ECG traces, recorded from wild-type (wt) and *Lmna*^+/-^ (heterozygous *Lmna*-KO) mice. Comparison of mean heart rate averaged over 1–2 min, P wave duration, P wave amplitude, PR interval, QRS duration and rate corrected QT interval (using the Fridericia formula) of wt (n = 8) or *Lmna*^+/-^ mice (n = 4). **b** Membrane capacitances, which serve as an index of cell size, are compared between atrial cardiomyocytes isolated from wt (n = 9 cells of N = 4 individuals) and *Lmna*^+/-^ (n/N = 18/5) mice. **c** Representative action potentials (APs), recorded from single atrial cardiomyocytes (CM) of both animal groups using the patch-clamp technique in current clamp configuration. Comparison of maximum upstroke velocity, AP amplitude, and AP duration at 50% or 90% of repolarization among wt (n/N = 9–10/4 animals) and *Lmna*^+/-^ (n/N = 10/4) mice. **d** Comparison of mean current densities of *I*_K,Peak_ (equivalent to the transient outward potassium current and the fast-activating slow-inactivating K^+^ current), quantified at +40 mV, mean current densities of the sustained potassium current component *I*_K,sus,_ quantified at +40 mV, *I*_K,Tail_ (equivalent to peak inward potassium current), quantified at −120 mV, and TASK-1 current-voltage (I-V) relationships, measured by application of the high-affinity TASK-1 inhibitor A293 (200 nM). Mean current *I*_TASK-1_ densities are quantified at +40 mV. Mean current values± SEM, derived from wt (n/N = 5–10/4) and *Lmna*^+/-^ (n/N = 4–8/4) CMs are displayed for each potassium current component. Data are presented as mean ± SEM. Dashed lines represent zero potential levels. Scalebars are provided as insets, p values were determined using two-tailed unpaired Student’s t-tests (*, p < 0.05; **, p < 0.01; ***, p < 0.001).

In isolated atrial CMs, membrane capacitance, as a surrogate for cell size, was significantly increased by 31.5 ± 9.16 % in *Lmna*^+/-^ mice, indicating atrial CMs hypertrophy (p = 0.0021; n/N = 27/9; Fig. 4b). However, current-clamp recordings of APs in these isolated cells revealed no differences in maximum upstroke velocity, AP amplitude, or APD_50_, while APD_90_ was shortened by 25.7 ± 8.6 % compared to wt (p = 0.012; n/N = 19-20/8; Fig. 4c). In summary, the triangulation of atrial APs was less pronounced in the murine laminopathy model compared to the *Rbm20*-R636Q model. A comprehensive analysis of the underlying ionic currents revealed no significant changes in *I*_K,Peak_, *I*_K,sus_, or *I*_K,tail_ currents. However, a striking 8.6-fold upregulation of the TASK-1 current (*I*_TASK-1_) was observed (p < 0.0001; n/N = 14/8; Fig. 4d), indicating a significant hallmark of atrial electrical remodeling in this *Lmna*^+/-^ mice.

### Transcriptomic characterization of the different murine disease models

To gain deeper insight into the molecular mechanisms underlying functional current changes, bulk RNA-seq analyses were conducted on atrial tissue samples from the various mouse models. A comparison of right atrial tissue between *Rbm20*-R636Q mice and wt controls (n = 5) identified 266 upregulated and 217 downregulated genes in the transgenic animals (Fig. 5a and c). Notably, several ion channel genes contributing to the altered ion currents were differentially expressed. For instance, the *Kcnj3* gene was significantly upregulated, alongside *Kcnd3* and *Scn5A*, while *Kcnj6* exhibited significant downregulation (Fig. 5a). Gene ontology (GO) analysis revealed that several of the top 20 categories were related to ion current regulation (Fig. 5b, red arrows) and extracellular matrix organization (Fig. 5b, green arrows). In contrast, a comparison with *Rbm20*-KO (n = 3–4) and *Lmna*^+/-^ (n = 3–4) mice revealed far fewer regulated genes (Fig. 5c). Although GO analysis in these models also pointed to dysregulation in ion transport processes, the extent was considerably lower. Moreover, transcriptomic profiling of atrial cardiomyopathy components highlighted that dysregulation of ion channel genes, along with genes implicated in myocardial fibrosis and inflammasome activation, was more prominent in *Rbm20*-R636Q mice compared to the KO or other dilated cardiomyopathy models like *Lmna*^+/-^(Fig. 5d, Suppl. Fig. 10).

**Figure 5:**
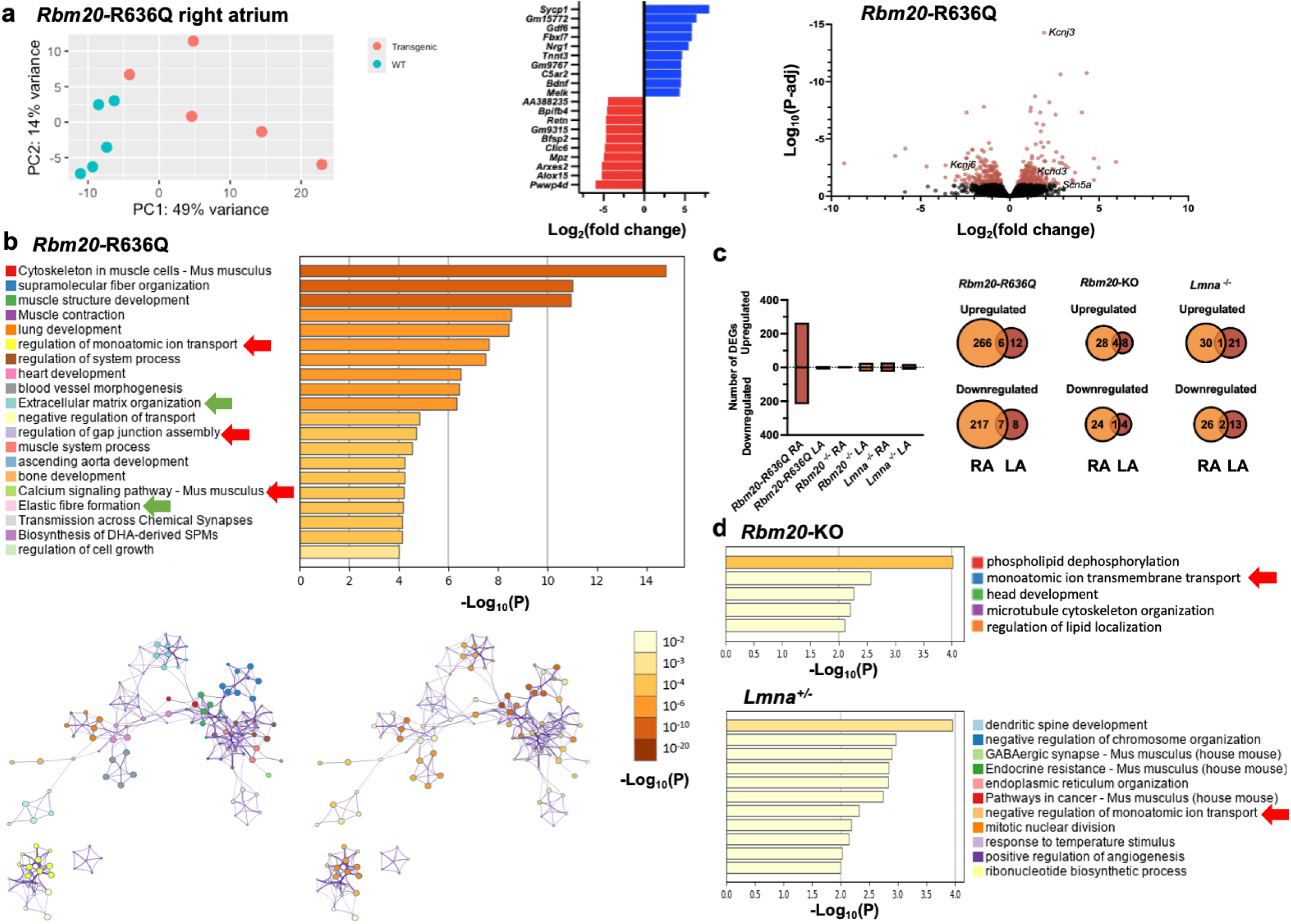
Transcriptomic characterization of Rbm20-related atrial cardiomyopathy. **a** *left:* PCA plot of bulk RNA-seq analysis from right atrial tissue samples of homozygous *Rbm20*-R636Q knock-in mice and corresponding wild-type (wt) controls (n = 5 each). *Center:* Top 10 upregulated and downregulated genes. *Right:* Volcano plot highlighting significantly regulated ion channel genes. **b** *Top:* Gene ontology (GO) analysis of differentially expressed genes in *Rbm20*-R636Q mice and corresponding wt controls. Pathways are shown with their corresponding −log_10_p values. GOs related to ion transport or extracellular matrix organization are highlighted with red and green arrows, respectively. *Bottom:* Graphical representation focusing on the interaction between different GOs. On the left, GOs are highlighted according to the color code shown at the top. The right figure shows the significance levels, color-coded as indicated. **c** Comparison of the number of differentially regulated genes (DEGs) in right atrial (RA) and left atrial (LA) tissue samples, derived from *Rbm20*-R636Q mice (n = 4–5), *Rbm20*-KO mice (n=3–4), and *Lmna*^+/-^ mice (n=3–4). *Bottom*: GO analysis of differentially expressed genes in *Rbm20*-KO and *Lmna*^+/-^ mice. GOs related to ion transport are highlighted with red arrows.

### Antiarrhythmic effects of SGLT-inhibitors in Rbm20-R636Q cardiomyocytes

Given that SGLT inhibitors have recently been incorporated into guideline-based HF therapy. As their beneficial effects on AtCM and AF incidence are being discussed, and peak sodium currents were shown to be significantly upregulated upon *Rmb20*-R636Q knock-in, the effects of different SGLT inhibitors on the cellular electrophysiology of *Rbm20*-R636Q CMs were evaluated.

AP inducibility, defined as the percentage of current injection pulses evoking an AP, out of 10 consecutive pulses applied at a rate of 0.5 Hz, was significantly reduced by increasing concentrations of the SGLT-1/SGLT-2 inhibitor sotagliflozin (25 µM: 25.7 ± 16.9 %; p = 0.18; n/N = 7/4 and 50 µM: 71.7 ± 16.5 %; p = 0.0072; n/N = 6/4; Fig. 6a–b). Additionally, a reduction in maximum upstroke velocity by 30.84 ± 13.8 % (p = 0.046; n/N = 5/4) was observed. The application of 25 µM sotagliflozin also counteracted the pathological APD_90_ shortening described in the *Rbm20*-R636Q model, with a significant increase in APD_90_ by 9.0 ± 4.0 % (p = 0.028; n/N = 5/4; Fig. 6b). Additionally a prolongation in APD_50_ by 11.8 ± 4.2 % was observed (p = 0.0032; n/N = 5/4; Fig. 6b).

**Figure 6:**
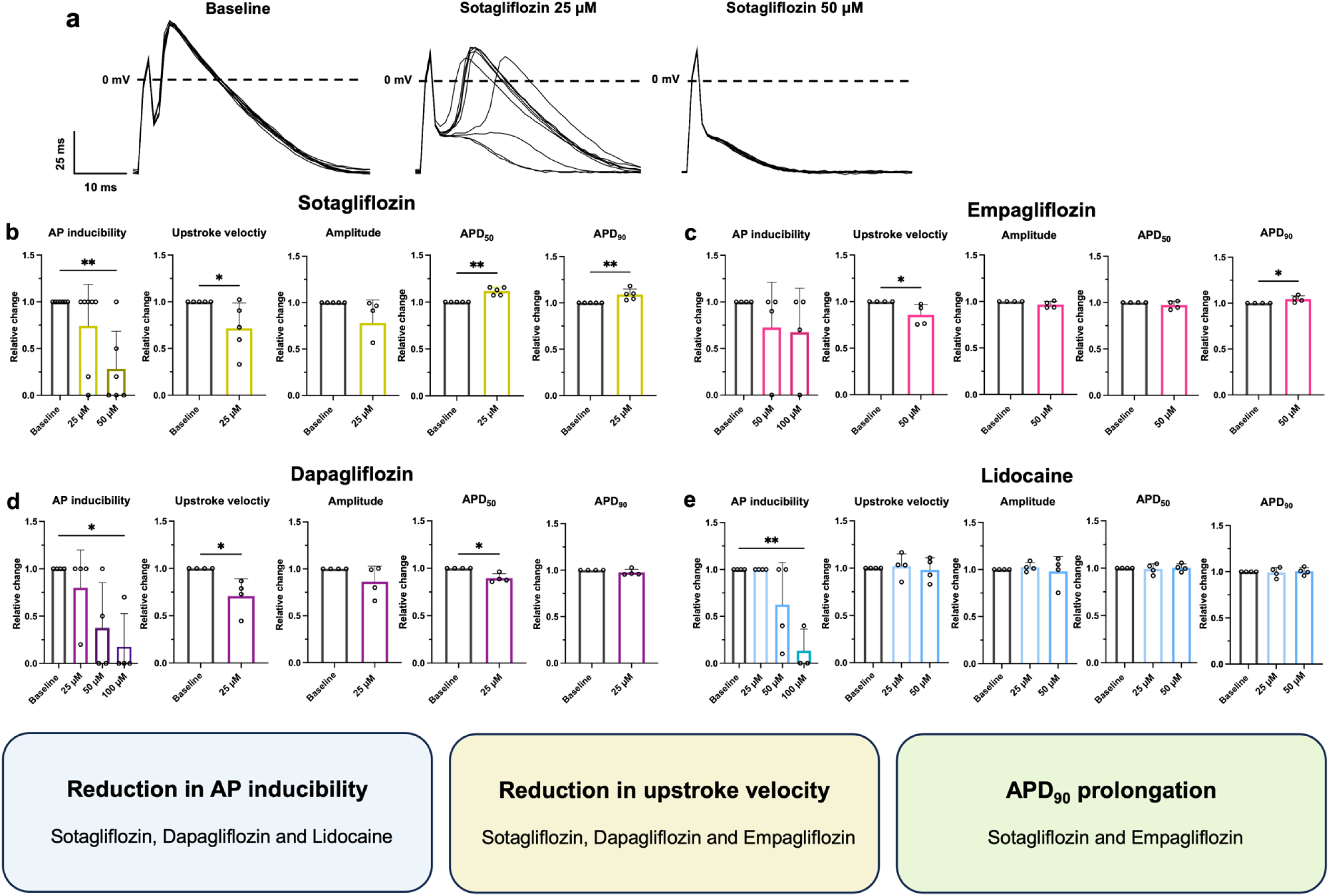
Electrophysiological effects of SGLT-inhibitors on action potential (AP) inducibility and shape in Rbm20-R636Q knock-in mice. **a** Current clamp recordings of 10 consecutive action potentials (APs) from a single atrial cardiomyocyte (CM), isolated from an homozygous *Rbm20*-R636Q mutant mouse, are shown under baseline conditions and after the administration of sotagliflozin at increasing concentrations (25 µM and 50 µM). **b–e** AP inducibility (i.e. the percentage of pulses evoking an AP out of 10 consecutive current pulses elicited at a rate of 0.5 Hz), maximum upstroke velocity, AP amplitude and AP duration at 50 % (APD_50_) 90 % (APD_90_) of repolarization, are compared at baseline and upon application of **(b)** sotagliflozin (25 µM and 50 µM; n = 4-7 cell derived from N = 4 animals), **(c)** empagliflozin (50 µM and 100µM; n/N = 4/4), **(d)** dapagliflozin (25 µM, 50 µM and 100 µM; n/N = 4/4) and **(e)** lidocaine (25 µM, 50 µM and 100 µM; n/N = 3-4/4) for reference as an *in class* sodium channel blocker. All data are provided as mean ± SEM. Dashed lines represent zero potential levels. Scalebars are provided as insets, p values were derived from two-tailed paired Student’s t-tests or, in case of three or more groups, one-way ANOVAs (*, p < 0.05; **, p < 0.01).

Weaker effects were observed for the SGLT2i empagliflozin, where a trend towards reduced AP inducibility was noted at concentrations of 50 µM and 100 µM, although this did not reach statistical significance (Fig. 5c). Empagliflozin at 50 µM mildly reduced the maximum upstroke velocity (14.4 ± 7.2 %; p = 0.084; n/N = 4/4; Fig. 5c), while the AP amplitude remained unchanged. However, a significant increase in APD_90_ by 6.9 ± 3.5 % was observed under empagliflozin treatment (p = 0.028; n/N = 4/4; Fig. 6c).

Similarly, the administration of the SGLT-2 inhibitor dapagliflozin resulted in a significant reduction in AP inducibility in *Rbm20*-R636Q cardiomyocytes (25 µM: 20.0 ± 10. %; p = 0.772, 50 µM: 62.5 ± 23.9 %; p = 0.073; n/N = 4/4; and 100 µM: 80.0 ± 17.5 %; p = 0.0170; n/N = 4/4; Fig. 5d). Dapagliflozin at 25 µM also significantly reduced the maximum upstroke velocity (29.3 ± 14.7 %; p = 0.050; n/N = 4/4) and APD_50_ (10.39 ± 4.6 %), while the AP amplitude and APD_90_ remained unchanged (Fig. 6d). In summary, these results indicate a direct anti-arrhythmic potential of those drugs on cellular level.

Since SGLT inhibitors have been shown to reduce peak sodium currents, it was important to determine whether the observed antiarrhythmic effects in Rbm20-R636Q cardiomyocytes (CMs) were due to inhibition of the upregulated sodium current. To investigate this, we also tested lidocaine, a well-known sodium channel blocker from Vaughan-Williams Class I. Lidocaine reduced action potential (AP) inducibility in a dose-dependent manner (100 µM: 86.7 ± 13.3 %; p = 0.0023; n/N = 3/3; Fig. 6e) but did not significantly affect maximum upstroke velocity, AP amplitude, APD50, or APD90. This suggests that SGLTi’s antiarrhythmic effects in Rbm20-R636Q CMs involve more than just inhibition of the peak sodium current. Furthermore, any potential impact of DMSO (used as a solvent for the drugs) was ruled out (Suppl. Fig. 7). These findings indicate that the antiarrhythmic effects of lidocaine occur at concentrations similar to those of the SGLT inhibitor.

## Discussion

For over a decade, it has been known that DCM patients harboring *RBM20* mutations exhibit a higher incidence of AF compared to patients with idiopathic forms of DCM^2^. Since then, GWAS have identified both *RBM20* and its splicing target *TTN* as two of the genes associated with AF in the general population^11,12,39^. More recently, *RBM20* mutations have been linked to the development of AF in both murine disease models and patients^16–18^. Despite these associations, the underlying proarrhythmogenic ionic alterations and the specific AtCM phenotype driving atrial arrhythmogenesis in RBM20-associated DCM remain, however, poorly understood.

The present study explores the role of loss of functional RBM20 mutations, particularly through the *Rbm20*-R636Q mutation, in the development of AtCM and its associated electrophysiological remodeling. The findings indicate that *Rbm20*-R636Q mice exhibit significant structural and functional atrial remodeling, including atrial enlargement, cardiomyocyte hypertrophy, and alterations in AP morphology, characterized by the triangulation of atrial APs which is pathognomonic for AF, APD_90_ shortening and an increased susceptibility to atrial arrhythmias. These results align with previous research linking RBM20 mutations to alterations in cellular electrophysiology of ventricular cardiomyocytes and arrhythmogenic outcomes^3,5,8,20^.

RBM20 functions as a splicing regulator for various essential cardiac genes involved in sarcomere organization and intracellular calcium regulation. This includes critical sarcomere-related genes such as *TTN*, *TPM1*, and *LDB3*, as well as genes responsible for calcium dynamics within the cell, like *CACNA1C*, *CAMK2D*, and *RYR2*^3^. Research conducted on CMs lacking *Rbm20* has shown elevated spontaneous Ca²⁺ release from the sarcoplasmic reticulum, which is attributed to an increased L-type Ca²⁺ current density^8^. This excess in intracellular Ca²⁺ and higher sarcoplasmic reticulum Ca²⁺ levels is likely driven by disrupted splicing of CaMKIIδ, leading to abnormal calcium handling^8^. Accordingly, in this work, a significant increase in L-type calcium currents was identified in isolated atrial CMs from *Rbm20*-R636Q mice, which contributes to the development of arrhythmogenesis.

As functional counterpart of this L-type calcium current density the outward transient potassium current was significantly decreased. In earlier publication the cytosolic voltage-gated potassium (K_v_) channel-interacting protein *KCNIP2* has been identified as a splicing target of RBM20^40^. The misplicing of this protein crucial for the function of the outward transient potassium channel group suggests that this remodeling is directly connected to the dysfunction of RBM20^3^. The resulting dysbalance of the outward transient potassium current with a strongly increased inward L-type calcium current density explains the promoted early repolarization time quantified by a prolonged APD_50_ in the *Rbm20*-R636Q model. In the other models characterizing the consequences of a loss of function on RBM20 included in this project this remodeling pattern did not appear which proposes this alteration to be specific for this model. Additionally, a significant upregulation of the atrial-specific TASK-1 potassium current density was noted, which is known to be a critical regulator of atrial APD and plays a role in the pathogenesis of AF^33,34,38,41^.

The upregulation of peak sodium currents in atrial *Rbm20*-R636Q cardiomyocytes observed in this study was correlated with increased transcriptional expression of the underlying *SCN5A* gene. Notably, elevated sodium currents have previously been associated with distinct subtypes of AtCM^27^. Interestingly, the peak sodium current density of atrial *Rbm20*-R636Q cardiomyocytes remains unchanged. This could be due to a significant downregulation of the voltage-gated outward transient potassium currents observed in these cardiomyocytes. As a result, sodium currents during phase I of the action potential experience less counter-regulation by early repolarizing potassium currents.

In summary, these changes in atrial ion currents provide a mechanistic explanation for the observed shortening of APD_90_ and AP triangulation, which predispose to atrial arrhythmogenesis by promoting reentry. Transcriptomic analysis further suggested extracellular matrix remodeling as an additional factor contributing to the observed pathophysiology.

To address the alterations in cardiac electrophysiology, SGLT inhibitors, widely used in HF treatment, demonstrated significant anti-arrhythmic potential by modulating AP morphology. While sotaglifozin, dapagliflozin, and empagliflozin all shared a class-I antiarrhythmic effect by inhibiting AP upstroke velocity, specific drug-dependent effects on cardiac electrophysiology were observed, suggesting differences in therapeutic potential and mechanisms of action. Notably, the class-III AP-prolonging effects of empagliflozin and sotagliflozin were especially relevant in counteracting the atrial AP shortening seen in RBM20 cardiomyopathy, warranting further investigation into the underlying potassium current inhibition.

Rare loss-of-function variants in RBM20 have further been linked to AF in two independent cohorts, including a Scandinavian cohort of early-onset AF patients and a separate cohort from the UK Biobank’s whole-exome sequencing data^16^. The study by Vad *et al.* could further provide experimental evidence that the loss of RBM20 leads to significant structural and transcriptional alterations in the atria. Their work highlighted changes in sarcomere organization and mitochondrial structure, as visualized by electron microscopy, and demonstrated that RBM20 loss impacts mitochondrial function. Just like the present study, these findings also suggest that RBM20 mutations disrupt not only the electrical but also the structural integrity of the atria, contributing to the development of AF^16^. In a subset of UK Biobank participants with cardiac magnetic resonance imaging data, RBM20 loss-of-function mutations were linked to abnormal atrial anatomy and function. This finding further supports the connection between RBM20 loss-of-function mutations and structural AtCM^16,18^.

Another study by Ihara *et al.* provided experimental evidence that the murine *Rbm20*-S367A mutation, located in close proximity to the mutation investigated in our study, also leads to a phenotype of both atrial and ventricular arrhythmias. Interestingly, similar to our findings, the *Rbm20*-KO genotype did not exhibit any spontaneous episodes of AF^17^.

Importantly, our observations were independent of the mice’s gender. This mirrors findings previously reported in the same mouse models regarding the ventricular phenotype and overall survival^5^.

Prolongation of the QTc interval in Rbm20-KO mice has been previously reported and was attributed to the disruption of RBM20-dependent alternative splicing^8^. Interestingly, this phenomenon was not observed in the present study. A possible explanation could be that, unlike in previous studies, the mice were not deeply sedated but were examined in a conscious state using the ecgTUNNEL system, which allows for potential autonomic nervous system influences on the QT interval.

HF is a well-established risk factor for the development and exacerbation of AF, with the two conditions often coexisting and interacting in a detrimental cycle. AF contributes to the worsening of HF by promoting irregular ventricular rates, leading to impaired cardiac output, increased myocardial oxygen demand, and further deterioration of ventricular function. Conversely, the structural and electrophysiological remodeling associated with HF, such as atrial dilation and fibrosis, predisposes patients to AF, which, in turn, accelerates disease progression and worsens overall outcomes. The presence of AF in HF patients has been associated with a higher incidence of hospitalizations, thromboembolic events, and mortality^9,42^. Therefore, early identification and treatment of AF in the absence of underlying cardiovascular disease is critical to prevent long-term complications. Equally or even more important is the management of AF in HF patients, where tailored interventions, including rate or rhythm control and anticoagulation, are essential to mitigate adverse outcomes. Timely and effective treatment of AF, particularly in the context of HF, can help break this vicious cycle and improve both arrhythmia control and HF prognosis. Therefore, recognizing and appropriately treating AF in RBM20-associated DCM patients is of utmost importance to prevent arrhythmia-related complications and improve overall disease outcomes.

Overall, the data of this study emphasize the central role of RBM20 in driving atrial electrical remodeling and highlight the therapeutic potential of SGLT inhibitors in modulating atrial electrophysiology and reducing AF susceptibility in the presence of *RBM20* mutations. This was further supported by transcriptomic analyses of atrial tissue samples from the transgenic animals, which revealed specific dysregulation patterns of key cardiac ion channel genes. However, gene dysregulations were not limited to ion channel genes; they also affected genes involved in the regulation of cardiac fibrosis and inflammasome activation. Finally, it was demonstrated that the dysregulation of these genes was more pronounced in tissue samples from *Rbm20*-R636Q mutants compared to *Rbm20*-KO or *Lmna* knockout samples.

### Potential limitations

It should be noted that the model used in this study carried a homozygous mutation, whereas most patients exhibit heterozygous variants. This highlights the need to investigate *RBM20*-related cardiomyopathy in a heterozygous model to gain further clinically relevant insights. While the Rbm20-R636Q mice used in this study were on a C57BL/6J background, the *Rbm20*-KO mice were bred on an FVB background. Differences in genetic backgrounds must be considered when interpreting electrophysiological and transcriptomic readout.

Regarding sex-specific differences, QRS prolongation was observed in male Rbm20-R636Q mice but not in females. However, no differences were noted in cellular electrophysiology between the sexes, which may explain this observation. Additionally, the limited number of male and female animals used in the study could have masked potential sex-dependent effects. A more robust analysis with a larger sample size is needed to more definitively assess sex-specific outcomes and better generalize the findings. Consistent with previous studies, the Rbm20-KO mice in this work did not exhibit spontaneous episodes of AF ^17^. The same applies to the laminopathy mouse model examined here. However, it should be noted that ECG measurements were taken only at discrete time points, and further studies using continuous telemetry ECG recordings would be necessary to definitively assess the AF burden in these animals. Moreover, inducing AF in mice is inherently challenging, and spontaneous AF is particularly rare, especially at such a young age. This factor may contribute to an underestimation of the AF burden in *Rbm20*-KO and laminopathy mice. Furthermore, the laminopathy model was established with a heterozygous KO which in general leads to a less pronounced phenotype.

### Conclusion

This study highlights the pivotal role of *RBM20* mutations, particularly *Rbm20*-R636Q, in driving atrial electrical remodeling and increasing susceptibility to AF. The observed alterations in AP duration, cardiomyocyte hypertrophy, and the upregulation of the atrial TASK-1 potassium current suggest that RBM20 dysfunction extends beyond its known association with DCM to significantly impact atrial electrophysiology. Furthermore, the demonstrated efficacy of SGLT inhibitors in modulating AP morphology and reducing arrhythmic potential in RBM20-mutant cardiomyocytes provides a promising therapeutic avenue. These findings underscore the potential of targeting atrial-specific ion channel dysregulation in the management of RBM20-related atrial cardiomyopathy and AF, offering new insights into both disease mechanisms and treatment strategies.

## Non-standard Abbreviations and Acronyms

AF: atrial fibrillation
AP: action potential
APD: action potential duration
AtCM: atrial cardiomyopathy
CaMKIIδ: Calcium/Calmodulin-dependent protein kinase II delta
CM: cardiomyocyte
DCM: dilated cardiomyopathy
DMSO: dimethyl sulfoxide
ECG: electrocardiogram
EGTA: ethylene glycol-bis(β-aminoethyl ether)-*N*,*N*,*N*′,*N′*-tetraacetic acid
ES: embryonic stem
FVB: Friend leukemia virus B strain
GEO: Gene Expression Omnibus
GWAS: genome-wide association studies
HEPES: 4-(2-hydroxyethyl)-1-piperazineethanesulfonic acid
KO: knockout
LA: left atrium
LV: left ventricle
MOPS: 3-morpholinopropane-1-sulfonic acid
RS: arginine-serine
SGLT: sodium-glucose co-transporter
SGLTi: sodium-glucose cotransporter inhibitors
SNP: single nucleotide polymorphism
TEA-Cl: tetraethylammonium chloride
TTN: titin
WT: wild-type

## Acknowledgments

We thank Anne Grube, Björn Rogatzki, Lisa Künstler, and Katrin Kupser, for their excellent technical support. The use of AI tools ChatGPT 4.o (OpenAI, San Francisco, California, USA) and DeepL (DeepL GmbH Cologne, Germany) to improve the readability of the manuscript is acknowledged, with the clarification that they were not employed for experimental design, data evaluation or interpretation.

## Supplemental Material

Supplemental Methods

Figures S1–S10

