## Supplement for "Disruption of RBM20 causes atrial electrophysiological disturbances"

**Short title:** *RBM20* mutation disrupts atrial electrophysiology

***Corresponding author:**

Prof. Dr. med. Constanze Schmidt, FESC, FEHRA

Department of Cardiology and Pneumology

University Medical Center Goettingen

Robert-Koch-Street 40

D-37075 Goettingen, Germany

Tel.: ++49 551 3967601

### Supplemental Methods:

#### *Animal ethics statement*

Animal experiments were conducted in compliance with guidelines set forth by the U.S. National Institutes of Health (NIH publication No. 86-23), the EU Directive 2010/63/EU, and German laws for animal protection. Approval was obtained from the local Animal Welfare Committee (Regierungspräsidium Karlsruhe, reference numbers G233/17, G225/20 and G-194/18). All animal care and procedures performed in this study, concerning *Rbm20*-R636Q knock-in mice, conformed to the EMBL guidelines for the Use of Animals in Experiments and were reviewed and approved by the Institutional Animal Care and Use Committee (IACUC).

#### *Mouse lines*

*Rbm20*-R636Q knock-in mice were generated via zygotic microinjection of recombinant Cas9 and *in vitro* reconstituted crRNA:trcrRNA complexes targeted to *Rbm20*, using a single-stranded donor DNA template as descried previously[20]. The B6C3F1 hybrid mouse strain was employed and subsequently backcrossed to C57BL/6J at the European Molecular Biology Laboratory (EMBL)[1]. *Rbm20* KO mice were originally generated at the Academic Medical Center, Amsterdam, using a conditional *Rbm20* allele produced by the International Knockout Mouse Consortium (University of California, Davis). A neomycin and LacZ cassette, along with loxP sites flanking exons 4 and 5, were inserted via homologous recombination in C57BL/6N embryonic stem (ES) cells. After pronuclear injection of these targeted ES cells into FVB blastocysts, Flp-mediated excision of the neomycin cassette was performed, followed by crossing with a CMV-Cre mouse line. To ensure a pure genetic background, *Rbm20* mice were backcrossed for at least six generations with FVB wild-type mice as previously described[8, 28]. The *Lmna* tm1.1Yxz/J line, was obtained from Jackson Laboratory and maintained on a C57BL/6J background as reported[29, 30].

#### *Animal handling, and anesthesia*

The animals were housed in individually ventilated plastic cages under controlled environmental conditions (temperature: 22 ± 2 °C, humidity 50 ± 20 %) with a 12-hour light/dark cycle. Mice were provided with ad libitum access to food and water. All experiments were conducted using male and female mice aged 15–17 weeks, with the specific number of animals used indicated in the corresponding figure legend. The patch clamp measurements for atrial action potentials and potassium currents were also replicated with 30 weeks old mice.

#### *Electrocardiography*

Electrocardiogram (ECG) recordings were performed on conscious mice using the non-invasive ecgTUNNEL system (emka Technologies, Paris, France), which allows for longitudinal monitoring in awake mice, eliminating the bias introduced by deep sedation, invasive procedures and pharmacological interference. Mice were gently restrained in the cylindrical tunnel to ensure proper paw contact with the system's silver electrode pads, which allowed for acquisition of 6-lead ECGs. Recordings were conducted at room temperature with the animal's spontaneous movements minimized to prevent artifacts. Each mouse was continuously monitored for approximately 5 min, during which a 6-lead ECG was recorded. From these sessions, around 3 min of high-quality, usable data were typically obtained. The ECG signals, including PR, QRS, QT, and RR intervals, were acquired via the ecgAUTO software, which facilitated high-throughput analysis using shape recognition techniques for arrhythmia detection. On average, over 100 heartbeats were analyzed per recording. The QT intervals were adjusted for heart rate using Fridericia's formula[31].

***Echocardiography***

Mice were sedated using a nose cone delivering 0.5 % to 2 % inhalative isoflurane, titrated to maintain a heart rate between 375 and 450 beats per minute (bpm). Time-gated, two-dimensional cine loops were acquired from parasternal long-axis, parasternal short-axis, and apical 4-chamber views using a Visualsonics Vevo 2100 system (Fujifilm, Tokio, Japan) and a MS550 transducer. To optimize visualization of the left atrium (LA), the standard murine parasternal long-axis window, displaying the left ventricular (LV) outflow tract, was adjusted slightly to the left. Two-dimensional left atrial size was measured using the Vevo LAB software (Fujifilm Tokio, Japan).

***Cardiomyocyte isolation***

Native murine atrial cardiomyocytes (CM) were isolated following euthanasia and rapid dissection of the mice. Left and right atrial tissue samples were immediately placed in a $Ca^{2+}$-free cardioplegic solution composed of (in mmol/L): 50 NaCl, 50 KCl, 6 $KH_2PO_4$, 25 $MgSO_4$, 250 taurine, 25 3-morpholinopropane-1-sulfonic acid (MOPS), 20 glucose, and 30 2,3-butanedione monoxime (pH 7.0) at 4–8 °C for transportation (15–60 min). Connective and fatty tissues were removed before the tissue was either flash-frozen in liquid nitrogen for subsequent transcriptomic analysis or maintained in cardioplegic solution and dissected into pieces of 0.5–1 $mm^3$ for enzymatic digestion. The tissue was rinsed in an ethylene glycol-bis(β-aminoethyl ether)-*N*,*N*,*N′*,*N′*-tetraacetic acid (EGTA)-containing buffer (in mmol/L: 137 NaCl, 5 $KH_2PO_4$, 1 $MgSO_4$, 5 HEPES, 10 glucose, 10 taurine, 0.2 EGTA; pH 7.4) for 5 min and subsequently digested in an EGTA- and $Ca^{2+}$-free solution with 200 U/mL collagenase type I (Sigma-Aldrich, Steinheim, Germany) and 5.4 U/mL protease type XXIV (Sigma-Aldrich, Steinheim, Germany) at 37 °C for 3-5 min. The digestion solution was oxygenated with 100 % $O_2$. The supernatant was discarded, and the tissue was stirred three times in protease-free solution for 2–5 min. Suspensions were centrifuged at 400 rpm for 2 min, and the resulting cell pellets were resuspended in a storage solution containing (in mmol/L): 10 EGTA, 25 glucose, 70 L-glutamic acid potassium salt monohydrate, 10 β-hydroxybutyrate, 20 KCl, 10 $KH_2PO_4$, 40 mannitol, 20 taurine, and 0.1 % albumin (pH 7.4). $Ca^{2+}$ was then gradually reintroduced to a final concentration of 2 mmol/L, and the cell suspension was maintained at room temperature.

***Cellular electrophysiology***

Consistently throughout all patch-clamp experiments, borosilicate glass patch pipettes (1B120F-4, World Precision Instruments, Berlin, Germany) with tip resistances of 6–13 MΩ were used. Recordings were made using an Axopatch 200B amplifier (Molecular Devices, San Jose, CA, USA) connected to an Axon Digidata 1550B interface (Molecular Devices, San Jose, CA, USA), with data acquisition and analysis performed using pCLAMP 11 software (Molecular Devices, San Jose, CA, USA). APs from both left and right atrial murine CMs were recorded under current clamp conditions at room temperature. The CMs were clamped with an average holding current density of -0.97 pA/pF, and APs were elicited by injecting current pulses of 250–800 pA (3 ms) at a frequency of 0.5 Hz. Sotagliflozin (Biozol, Eching, Germany), dapagliflozin (Biozol, Eching, Germany), empagliflozin (Biozol, Eching, Germany) and lidocaine (Sigma Aldrich) were dissolved in dimethyl sulfoxide (DMSO) to prepare 200 mM stock solutions and stored at -20°C. After stabilization of the APs, sotagliflozin, empagliflozin, and dapagliflozin were introduced into the chamber at the specified concentrations. A293 was synthesized by Enamine Ltd. (Kyiv, Ukraine) and dissolved in DMSO at a concentration of 10 mM. The final DMSO concentration in the chamber was maintained at ≤0.04 %, ensuring it did not significantly influence AP parameters in atrial CMs, in accordance with previous studies[27]. For AP recordings, the internal (pipette) solution contained (in mmol/L) 134 potassium D-gluconate,

6 NaCl, 1 MgATP, 10 HEPES (pH 7.2) and CMs were placed in an external solution containing (in mmol/L) 137 NaCl, 5.4 KCl, 2 $CaCl_2$, 1 $MgSO_4$, 10 HEPES, and 10 glucose (pH 7.3). Clamped resting membrane potentials did not differ among the experimental groups (Suppl. Fig. S1).

L-type calcium currents were obtained by voltage clamp recordings, employing a whole-cell ruptured-patch configuration. From a holding potential of -80 mV, a 400 ms ramp pulse to -40 mV was applied to inactivate fast sodium ($Na^+$) currents, followed by 100 ms test pulses with voltage steps incrementing by 10 mV, ranging from -60 mV to +60 mV, with an additional step to +10 mV. These pulses were delivered at a frequency of 0.5 Hz. The internal solution contained (in mmol/L) 0.02 EGTA, 0.1 GTP-Tris, 10 HEPES, 92 K-aspartate, 48 KCl, 1 Mg-ATP, and 4 $Na_2$-ATP (pH=7.2) and the external bath solution consisted of (in mmol/L) 2 $CaCl_2$, 10 glucose, 10 HEPES, 4 KCl, 1 $MgCl_2$, 140 NaCl, and 2 probenecid (pH = 7.4). For voltage-clamp experiments, $K^+$ currents were blocked by applying 5 mM 4-aminopyridine and 0.1 mM $BaCl_2$ to the bath solution.

To assess voltage-dependent sodium currents, cardiac myocytes (CMs) were voltage-clamped (whole-cell ruptured-patch configuration) at a holding potential of -100 mV. Sodium currents were elicited by applying successive 10 mV voltage steps (500 ms), ranging from -100 mV to +20 mV. The internal solution contained (in mmol/L) 100 potassium L-aspartate, 20 KCl, 2 $MgCl_2$, 1 $CaCl_2$, 2 $Na_2$ATP, 10 EGTA, and 10 HEPES (pH 7.2) while the external bath solution consisted of (in mmol/L) 10 NaCl, 2 $CaCl_2$, 3 MgATP, 135 CsCl, 2 TEA-Cl, 5 EGTA, and 0.2 HEPES (pH 7.2).

Potassium currents were measured using voltage clamp recordings in a whole-cell ruptured-patch configuration with voltage pulses applied in 10 mV increments from −60 mV to +60 mV, originating from a holding potential of −80 mV. A 10 ms prepulse to −40 mV was used to inactivate sodium channels. Each depolarizing step of 500 ms was followed by a 100 ms pulse to −120 mV. This protocol allows for the assessment of the inward transient $K^+$ current (at the onset of the voltage pulse), of sustained $K^+$ currents (at the end of the voltage pulse), and the inward rectifying potassium channels (during the hyperpolarizing second pulse) as described previously[32]. Furthermore, TASK-1 potassium channels were isolated from outward potassium currents using the high-affinity TASK-1 inhibitor A293 at a concentration of 200 nM as previously described[33, 34]. The internal solution contained (in mmol/L) 5 NaCl, 120 KCl, 2.5 MgATP, 1 EGTA, and 5 HEPES (pH 7.2) and the external bath solution consisted of (in mmol/L) 136 NaCl, 5.4 KCl, 1 $CaCl_2$, 1 $MgCl_2$, 5 HEPES, 0.33 $NaH_2PO_4$, and 10 glucose (pH 7.4).

***RNAseq***

Murine left and right atrial tissue samples, collected as described above, were flash frozen in liquid nitrogen and stored at -80 °C. Tissue samples were homogenized using a TissueRuptor (QIAGEN, Hilden, Germany) system and a total RNA fraction was isolated using TRIzol Reagent (Invitrogen, Thermo Fisher Scientific, Waltham, MA, USA) according to the manufacturer's instructions. 500 ng of total RNA was subjected to poly(A)-enriched bulk RNAseq (Illumina NextSeq 2000). Reads were aligned to GRCm39 using STAR (version 2.7.11b)[35]. Read count files were generated with featureCounts (version 2.0.3)[36] for each gene. For differential gene expression analysis, DESeq2 (version 1.44.0) was used, and pairwise comparisons between genotypes were performed[37]. Adjusted p values were calculated using the Benjamin & Hochberg method. A gene was considered differentially expressed if the $\log_2$ of its expression fold change was greater than 0.5, and if the adjusted p value was less than 0.01.

#### *Statistical analysis*

Statistical analysis and data visualization were conducted with Prism 10 (GraphPad Software, La Jolla, CA, USA). The data are presented as mean ± standard error of the mean (SEM). Paired or unpaired two-tailed Student's t-tests, one sample Student's t-tests as well as one-way or two-way ANOVA with Tukey's multiple comparison posttest or log-rank test were used to determine statistical significance. The name of the test, p value and number of biological replicates are indicated in each figure legend. In case of small sample sizes normality was assessed using Shapiro-Wilk tests. A p value of less than 0.05 was considered statistically significant. If the null hypothesis of equal means is rejected at the 0.05 significance level, pairwise comparisons among groups were conducted and the probability values were adjusted for multiple comparisons using the Bonferroni correction.

#### *Data availability statement*

The main data is available in the main text or the supplementary materials. Bulk RNA-seq data will be uploaded to NCBI GEO upon publication of the manuscript. All further data are available from the corresponding author upon reasonable request.

### Supplemental Figures:

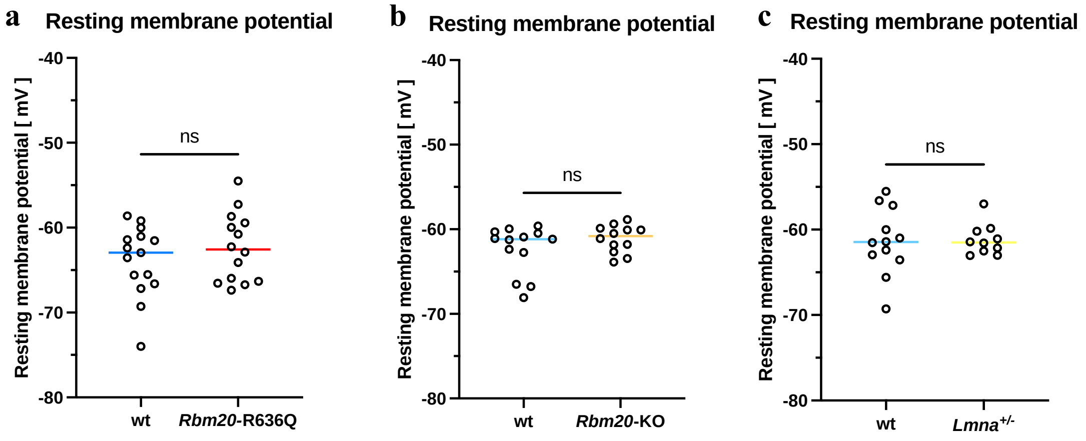


***Figure S1: Comparison of clamped resting membrane potentials among all experimental groups***

Clamped resting membrane potential during action potential recordings, depicted in Fig. 2, 3 and 4, respectively. **a** (wt n/N = 15/6, *Rbm20*-R636Q n = 14/5), **b** (wt n/N = 12/4, *Lmna*$^{+/-}$ n/N = 10/4), **c** (wt n/N = 13/4, *Rbm20*-KO n/N = 12/4). P-values were derived from unpaired Student's t-tests. ns, not significant.

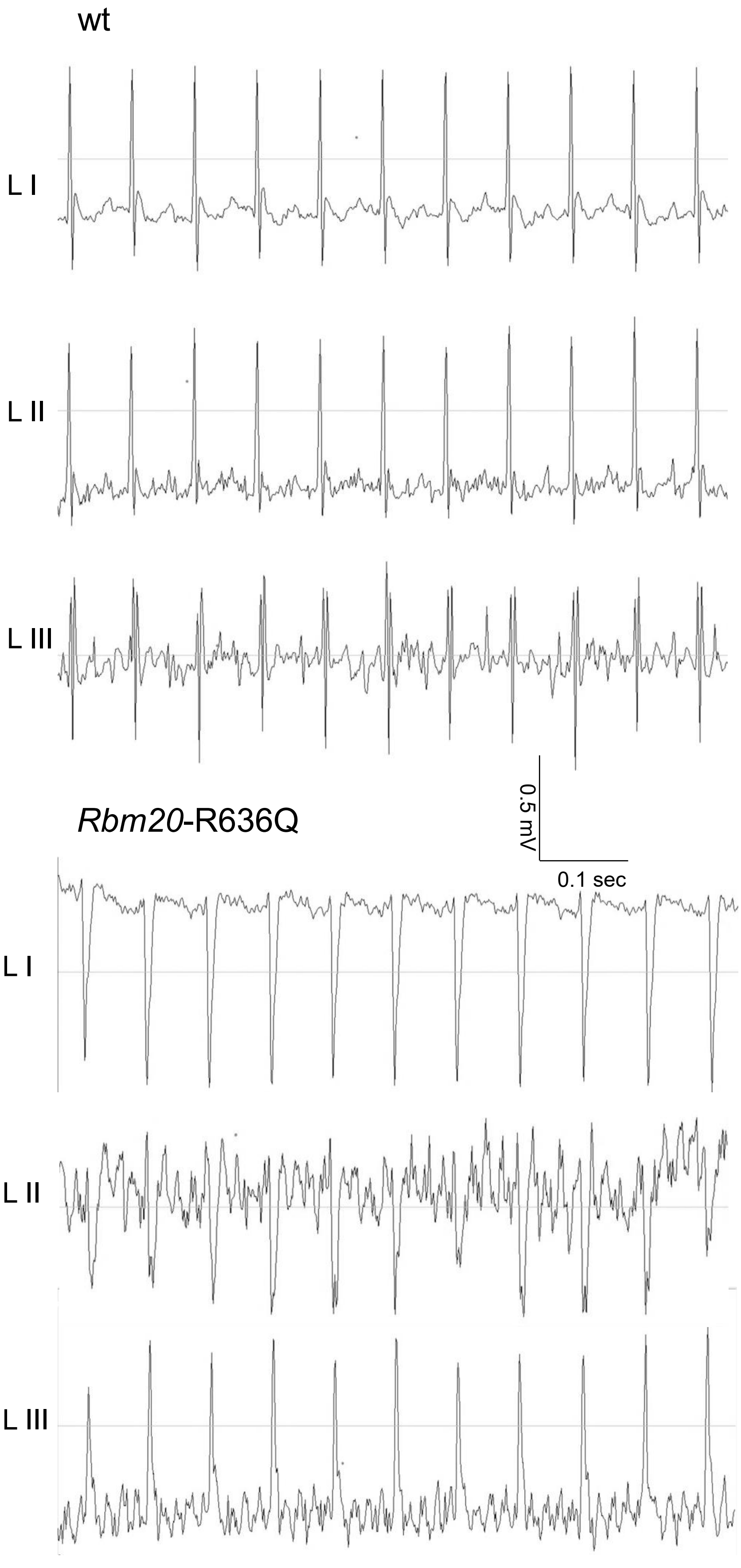
wt
L I
L II
L III
0.5 mV
0.1 sec
Rbm20-R636Q
L I
L II
L III

***Figure S2: Representative ECGs***

Representative surface ECG recordings (60 s) of a wild type and *Rbm20*-R636Q mice. Limb leads according to Einthoven I–III as indicated.

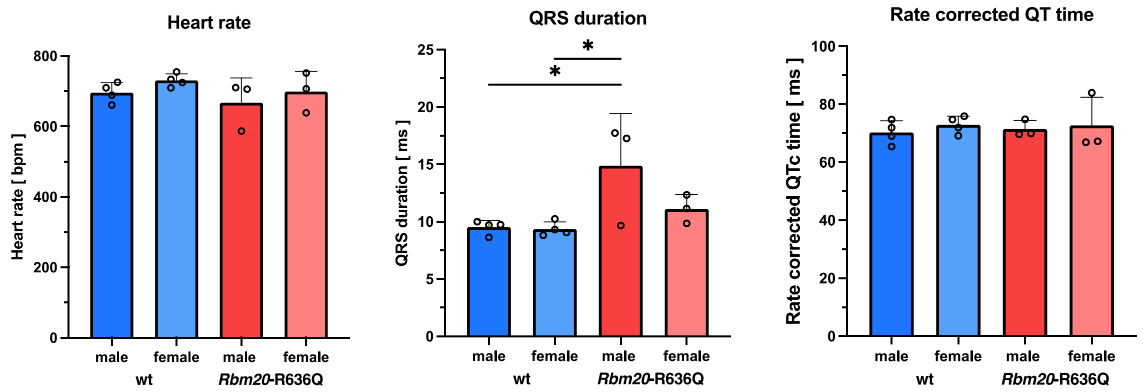


***Figure S3: Differences in ventricular electrophysiology in Rbm20-R636Q between genders in non-invasive ECGs***

Analysis of non-invasive ECG measurements (wt male n/N = 4/3, female n/N = 4/4; *Rbm20*-R636Q male n/N = 3/3, female n/N = 3/3) with corresponding quantification of heart rate, QRS duration and rate corrected QT time using the Fridericia formula, respectively. All data are shown as mean ± SEM. P-values were derived from ordinary one-way analysis of variance (ANOVA).

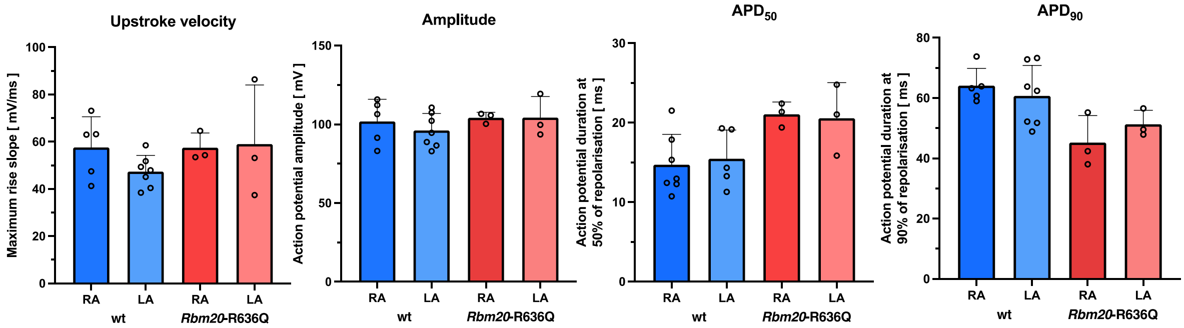


***Figure S4: No differences in atrial action potential morphology between left and right atria cardiomyocytes***

Analysis of atrial action potentials (wt RA n/N = 5/3, LA n/N = 7/3; *Rbm20*-R636Q RA n/N = 3/3, LA n/N = 3/3) with corresponding quantification of maximum upstroke velocity, AP amplitude and AP duration at 50% ($APD_{50}$) and 90% ($APD_{90}$) of repolarization, respectively. Data are shown as mean ± SEM. P-values were derived from ordinary one-way analysis of variance (ANOVA).

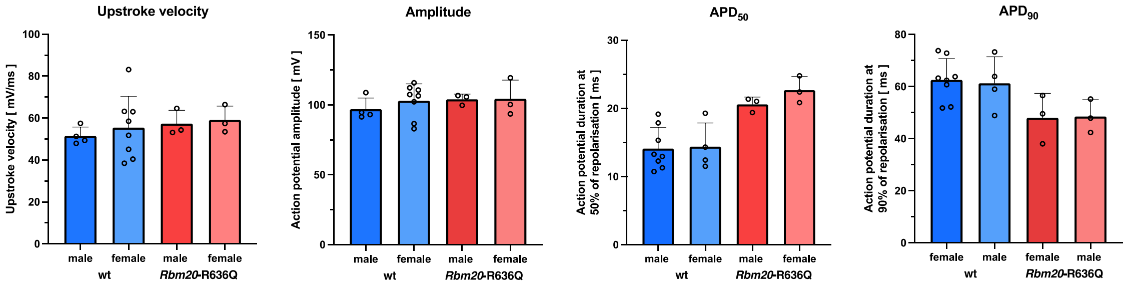


***Figure S5: No differences in atrial action potential shape between male and female mice***

Analysis of atrial action potentials (wt male n/N = 4/3, female n/N = 8/4; *Rbm20*-R636Q male n/N = 3/3, female n/N = 3/3) with corresponding quantification of maximum upstroke velocity, AP amplitude and AP duration at 50% ($APD_{50}$) and 90% ($APD_{90}$) of repolarization, respectively. Data are depicted as mean ± SEM. P-values were derived from ordinary one-way analysis of variance (ANOVA).

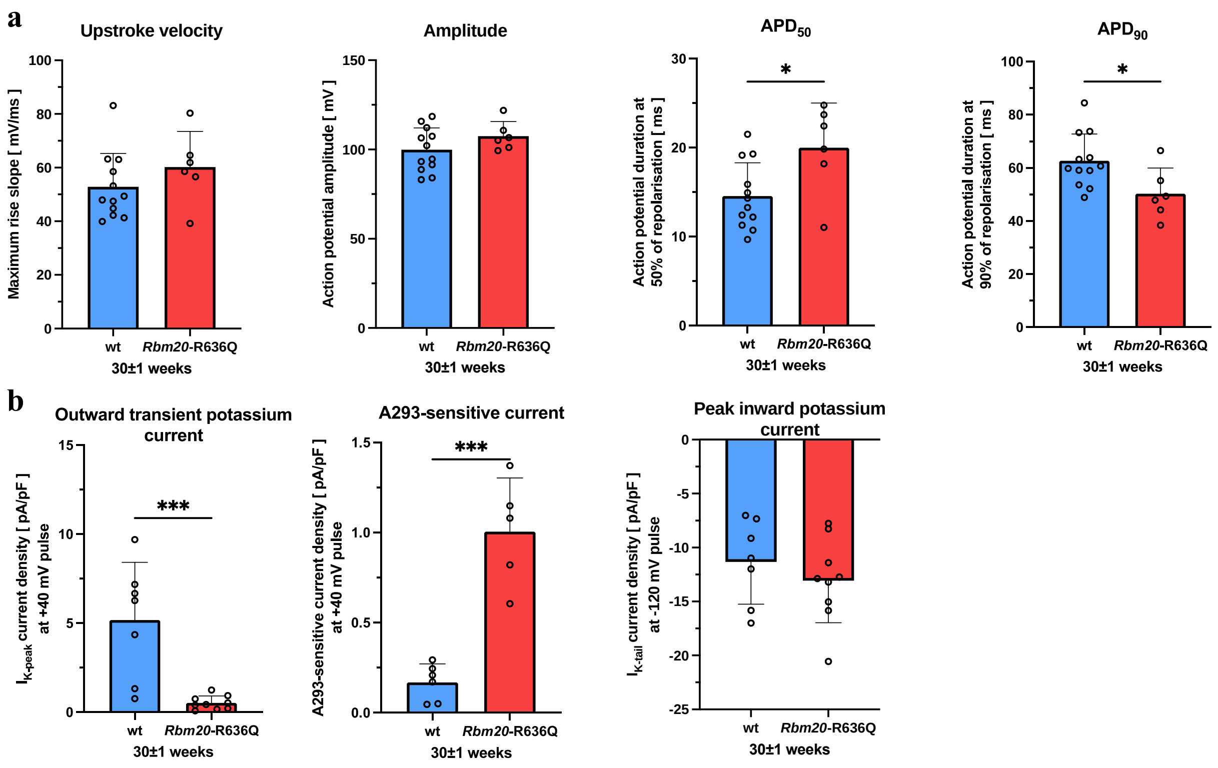


***Figure S6: Similar remodeling in 30-week-old mice***

**a** Representative atrial action potentials (AP) of isolated cardiomyocytes of animal groups (wt n/N = 12/4, *Rbm20*-R636Q n/N = 6/3) as indicated with corresponding quantification of max. upstroke velocity, AP amplitude and duration at 50% ($APD_{50}$) and 90% ($APD_{90}$) of repolarization **b** Potassium currents corresponding to the measurements in Fig. 2 analyzed as of $I_{K,Peak}$ (wt n/N = 7/4, *Rbm20*-R636Q n/N = 9/3) caused by the voltage protocol after the step to -40 mV and quantified at the +20 mV pulse, to $I_{TASK-1}$ (wt n/N = 7/4, *Rbm20*-R636Q n/N = 9/3) analyzed at the end the voltage step as difference between baseline and 200 nM A293 and quantified after the +40 mV pulse and $I_{K,Tail}$ (wt n/N = 7/4, *Rbm20*-R636Q n/N = 9/3) analyzed at the end -120 mV step and quantified after the +60 mV pulse. All data are shown as mean ± SEM. P-values were derived from unpaired Student’s t-tests (*, p < 0.05; **, p < 0.01; ***, p < 0.001).

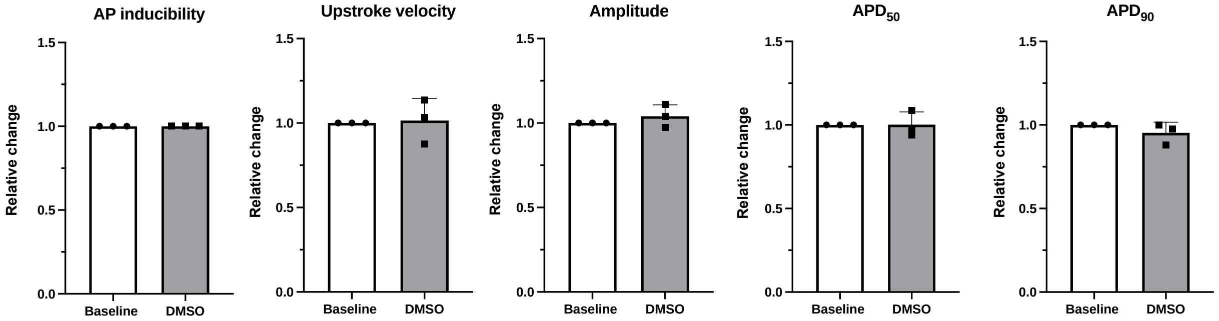


***Figure S7: DMSO has no effect on atrial action potential***

No alterations by DMSO in AP inducibility (percentage of pulses evoking an AP, out of 10 current pulses elicited at a rate of 0.5 Hz), maximum upstroke velocity, AP amplitude and AP duration at 50 % ($APD_{50}$) and 90% ($APD_{90}$) of repolarization, at baseline and under 7 min drug application (DMSO 1:800 corresponding to maximum concentrations used during drug application). *Rbm20*-R636Q n/N = 3/3. All data are shown as mean ± SEM. P-values were derived from paired Student's t-tests.

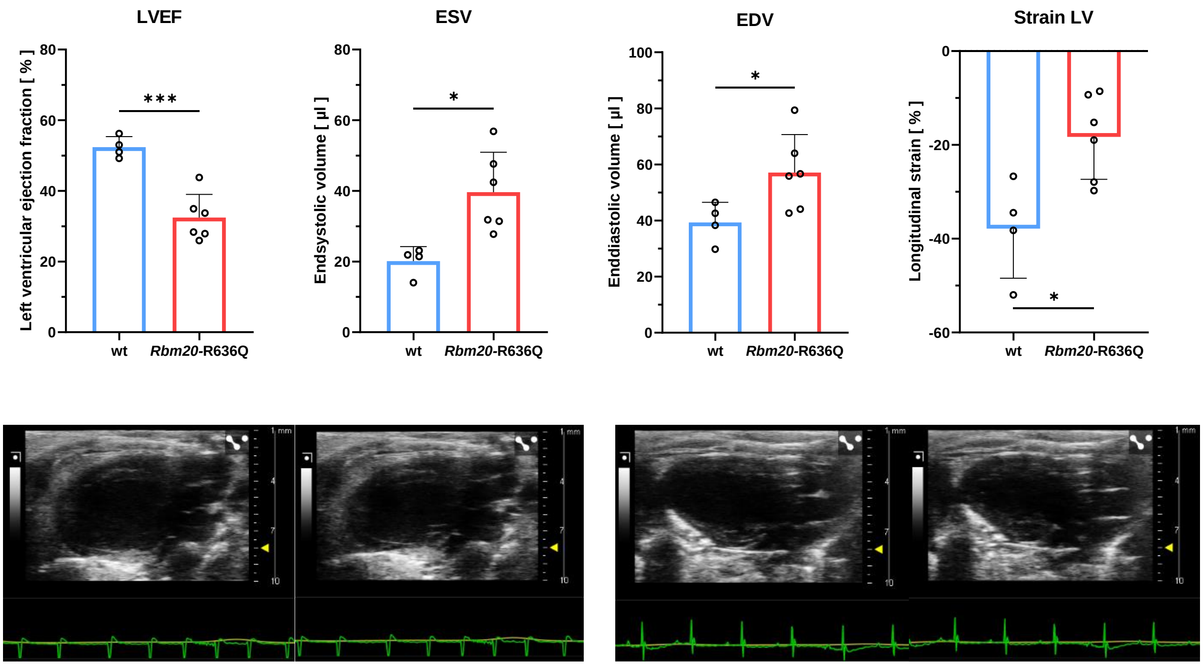


*Rbm20*-R636Q diastolic *Rbm20*-R636Q systolic wt diastolic wt systolic

***Figure S8: Echocardiographic analysis of ventricular ejection fraction and longitudinal strain***

*Top:* Analysis of left ventricular ejection (LVEF), endsystolic volume (ESV), enddiastolic volume (EDV) and longitudinal strain of *Rbm20*-R636Q and wild-type (WT) mice shown in Fig. 1 (WT n = 4, *Rbm20*-R636Q n = 6). *Bottom:* Corresponding representative echocardiographic images, obtained from parasternal long-axis view with corresponding ECG. All data are shown as mean ± SEM. P-values were derived from paired Student's t-tests (*, $p < 0.05$; ***, $p < 0.001$).

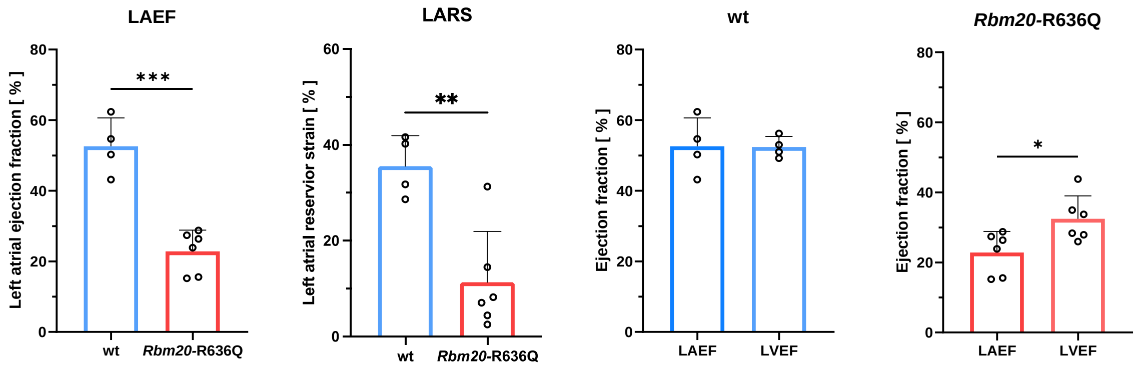


***Figure S9: Echocardiographic analysis of atrial function***

*Left:* Analysis of left atrial ejection fraction (LAEF) and left atrial reservoir strain of *Rbm20*-R636Q and wild-type (WT) mice shown in Fig. 1 (WT n = 4, *Rbm20*-R636Q n = 6). *Right:* Comparison of left atrial and left ventricular ejection fraction in wild-type and *Rbm20*-R636Q mice already analyzed in Fig.1 and S09. All data are given as mean ± SEM. P-values were derived from paired Student's t-tests (*, $p < 0.05$; **, $p < 0.01$; ***, $p < 0.001$).

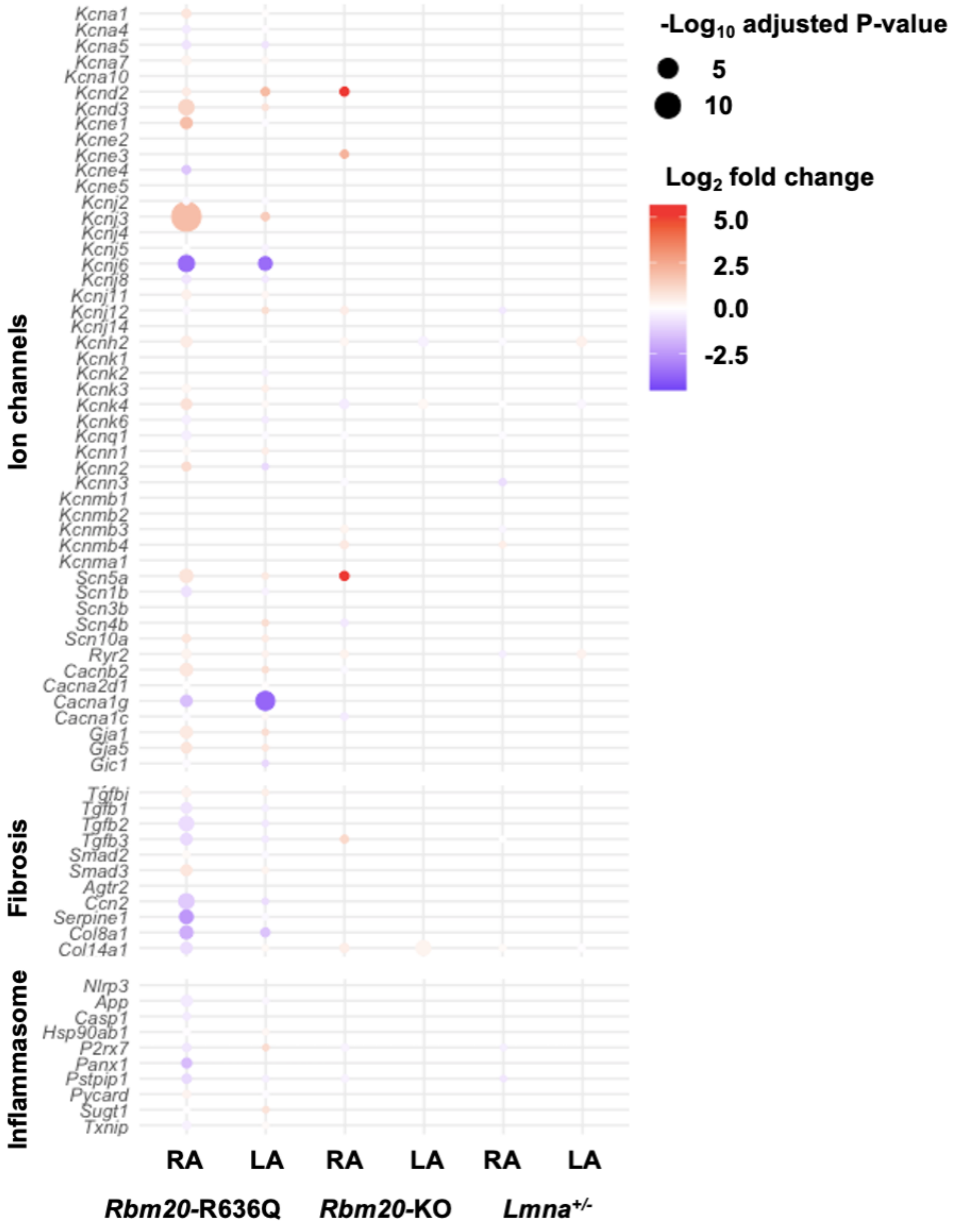


***Figure S10: Transcriptomic analysis of atrial cardiomyopathy components in different murine models of dilated cardiomyopathy (DCM)***

Dot plot visualizing the $\log_2$ fold change (color) and $-\log_{10}$ adjusted p values (dot size) of various genes contributing to ion channel remodeling, fibrosis formation, and inflammasome activation in atrial cardiomyopathy. Comparisons are shown for right atrial (RA) and left atrial (LA) tissue samples from *Rbm20*-R636Q mice (n = 4–5), *Rbm20*-knockout mice (n = 3–4), and *Lmna*$^{+/-}$ mice (n = 3–4) against their respective control animals.
